# Control of recollection by slow gamma dominating mid-frequency gamma in hippocampus CA1

**DOI:** 10.1101/152488

**Authors:** Dino Dvorak, Basma Radwan, Fraser T. Sparks, Zoe Nicole Talbot, André A. Fenton

**Affiliations:** Center for Neural Science, New York University, New York, 10003, USA; School of Medicine, New York University, New York, NY 10016, USA; Neuroscience Institute at the New York University Langone Medical Center, New York, NY, 10016, USA; Department of Physiology and Pharmacology, The Robert F. Furchgott Center for Neural & Behavioral Science, State University of New York, Downstate Medical Center, Brooklyn, New York, 11203, USA

## Abstract

Behavior is used to assess memory and cognitive deficits in animals like Fmrl-null mice that model Fragile X Syndrome, but behavior is a proxy for unknown neural events that define cognitive variables like recollection. We identified an electrophysiological signature of recollection in mouse dorsal CA1 hippocampus. During a shocked-place avoidance task, slow gamma (SG: 30-50 Hz) dominates mid-frequency gamma (MG: 70-90 Hz) oscillations 2-3 seconds before successful avoidance, but not failures. Wild-type but not Fmrl-null mice rapidly adapt to relocating the shock; concurrently, SG/MG maxima (SG_dominance_) decrease in wild-type but not in cognitively inflexible Fmrl-null mice. During SG_dominance_, putative pyramidal cell ensembles represent distant locations; during place avoidance, these are avoided places. During shock relocation, wild-type ensembles represent distant locations near the currently-correct shock zone but Fmrl-null ensembles represent the formerly-correct zone. These findings indicate that recollection occurs when CA1 slow gamma dominates mid-frequency gamma, and that accurate recollection of inappropriate memories explains Fmrl-null cognitive inflexibility.

## INTRODUCTION

The hippocampus is crucial for both learning and remembering information, especially about space [1], and because the same place-representing neurons participate in both processes [2-7], it is unknown what neural events control whether hippocampal neurons are encoding current experience or recollecting information from memory [8]. A prominent “communication-through-coherence” [9-12] or “routing-by-synchrony” hypothesis asserts that activity in CA1 switches between an information acquiring mode associated with mid-frequency 60-90 Hz gamma oscillations that synchronize hippocampus output with neocortical input and a separate long-term memory recollection mode associated with slow 30-60 Hz gamma oscillations that synchronize CA1 output with intrahippocampal CA3→CA1 inputs [12, 13]. Gamma oscillations are generated by local interneurons [14-17] and the local CA1 GABAergic currents that underlie gamma oscillations are effectively driven by tonic excitation, as described by pyramidal interneuron network gamma (PING) models of gamma generation [18-20]. Furthermore, tonic inputs to PING as well as interneuron network gamma (ING) models can locally generate distinct lower and higher frequency gamma oscillations by local competition between distinct interneuron populations with correspondingly long and short lasting post-synaptic inhibition [21]. Because CA1 receives two anatomically distinct inputs [22, 23] and each mediates both dendritic excitation and feedforward inhibition [24], routing-by-synchrony hypotheses predict that during long-term memory recall the CA3-associated slow gamma input will outcompete the entorhinal cortex-associated mid-frequency gamma input for control of CA1 output.

We test this prediction and find in freely-behaving mice solving a place task, that slow and mid-frequency gamma oscillations are concurrent in mouse CA1, but a transient dominance of slow gamma oscillations over mid-frequency gamma oscillations signals recollection. This slow gamma dominance lasts several hundred milliseconds and occurs on average every ˜9 seconds, both when mice are active or still. Increased and decreased rates of slow gamma dominance predicts accurate, failed and changed place memory in wild-type mice, as well as cognitive inflexibility in a Fmr1-null mutant mouse model of Fragile X Syndrome intellectual disability, associated with high prevalence of autism. During slow gamma dominance, putative pyramidal cell ensemble discharge represents distant locations, and during place avoidance tasks these distant locations are the vicinity of the shock zone that the mouse learned to avoid. However, when Fmr1-null mice express cognitive inflexibility by continuing to avoid the formerly correct and now incorrect place, these slow gamma dominance events are excessive and predictive of putative pyramidal cell representations of formerly-correct shock-zone location memories. Because gamma oscillations are generated by local inhibitory synapses, and consistent with theory [21], these results point to local biases in competing gamma-generating inhibitory events as the potential origin of distinct information and long-term memory processing modes, such as recollection.

## RESULTS

### Identifying recollection events prior to active avoidance

We began by identifying when mice were likely to recall the location of shock during training in variants of the active place avoidance task [Fig. 1A; 25]. Periods of stillness, when the mouse is passively carried towards the shock zone, are interrupted by active avoidances (Fig 1B), indicating successful recollection of the shock location and identifying times with a high likelihood of recollection (Fig 1).

**Fig 1.**
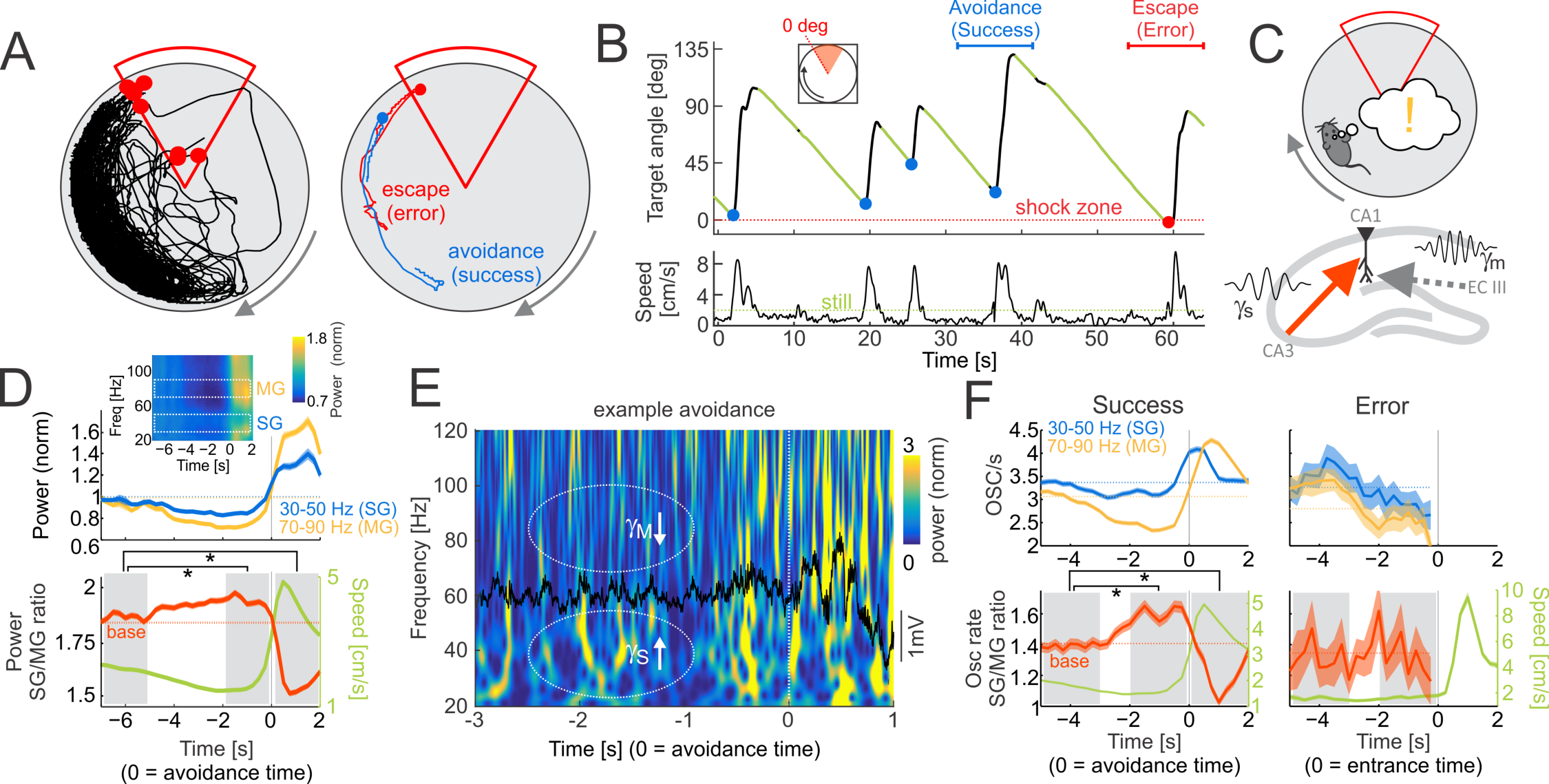
Slow gamma dominates mid-frequency gamma prior to successful place avoidance. (A) Left: typical 30-min path during the third active place avoidance training session. Shocks are shown as red dots. Right: Example of avoidance (success; blue line) and escape after receiving a shock (error; red line). (B) Top: time profile of the angular distance to the leading edge of the shock zone showing a typical saw-tooth avoidance pattern during ˜60 s. Periods of stillness (green), when the mouse is passively carried towards the shock zone are interrupted by active avoidances (blue dots). Entrance to the shock zone is marked as a red dot. The horizontal blue and red lines mark time intervals of the example avoidance and escape from panel A, right. The red dotted line marks the leading edge of the shock zone. Bottom: speed profile during the same ˜60-s interval. The stillness threshold is shown as a green dotted line at 2 cm/s. (C) Schematic depiction of the working hypothesis – as the mouse approaches the shock zone (top), slow gamma driven by CA3 inputs transiently dominates mid-frequency gamma driven by ECIII inputs causing recollection of the shock zone location. (D) Top: average power of slow gamma (blue; 30-50 Hz) and mid-frequency gamma (yellow; 70-90 Hz) in the LFP around the time of avoidance initiation (T=0). Mean powers are displayed as dotted lines. Inset shows average of normalized power across 20-120 Hz around avoidance initiation. Representative slow and mid-frequency gamma bands are marked by white rectangles. Bottom: the average ratio of slow to mid-frequency gamma power (red line) around avoidance initiation. The mean power ratio is shown as a dotted line. The corresponding average speed profile is shown in green. Data are represented as average ± SEM. Gray boxes represent time intervals for statistical comparisons, *p < 0.05 relative to baseline (-7..-5 s). (E) The time-frequency representation of a 4-s example LFP (overlayed in black) around the initiation of an avoidance start (T=0 marks avoidance initiation). Notice the relative reduction in number of mid-frequency gamma (70-90 Hz) oscillatory events relative to slow gamma (30-50 Hz) events prior to the avoidance (T ˜ -2 s) compared to times during the active avoidance (T > 0 s). (F) Left, top: average event rates for slow gamma (blue; 3050 Hz) and mid-frequency gamma (yellow; 70-90 Hz) oscillations around the time of avoidance initiation (T=0). Mean rates are displayed as dotted lines. Left, bottom: the average ratio of slow to mid-frequency gamma event rates (red line) around avoidance initiation. The mean ratio is shown as a dotted line. The corresponding average speed profile is shown in green. Right: same as F, left but for Avoidance errors. Data are represented as average ± SEM. Gray boxes represent time intervals for statistical comparisons, *p < 0.05 relative to baseline (-5..-3 s).

The routing-by-synchrony hypothesis [12, 13] predicts that CA3-driven slow gamma oscillations will transiently dominate neocortex-driven mid-frequency gamma oscillations when the mouse is recollecting the shock zone location (Fig 1C). Concurrent local field potentials (LFPs), reflecting synchronous synaptic activity within the dorsal hippocampus, were recorded at the perisomatic region of CA1 and examined during these behavioral segments with high likelihood of recollection. The LFP state was mostly in theta, although somewhat lower amplitude during stillness (Supplementary information Fig S1A), as is typical for spatially alert stillness [26]. At *stratum pyramidale*, slow and mid-frequency gamma power could be separated by their different phase relationships to theta but less so by their frequency content during both stillness and active locomotion (Supplementary information Fig S2C, D). Importantly, the rate of sharp-wave associated ripples during these pre-avoidance periods of stillness was no different than the overall stillness ripple rate (Supplementary information Fig S1C).

It was reported that theta oscillations in the *stratum pyramidale* LFP of the free-behaving rat are predominantly concurrent with either 25-50 Hz CA3-associated gamma or 65-140 Hz entorhinal cortex layer 3 (ECIII) gamma oscillations, but rarely both, and the slower gamma tends to occur at an earlier theta phase than the faster gamma [12]. In contrast to those recordings in the rat, we find that in the mouse, both slow and mid-frequency gamma oscillations frequently occur within single theta oscillations in the *stratum pyramidale* LFP (see Supplementary information Fig S2E and Fig S3D). It is only after selecting oscillations with the largest power that single theta cycles can be shown to be dominated by either slow or mid-frequency gamma oscillations, but this is likely an artifact of rejecting most oscillations since only a single supra-threshold gamma oscillation occurs within a single theta cycle when the threshold is > 2 S.D. (Supplementary information Fig S3D). Furthermore, we also find in the mouse that slow gamma oscillations occur close to the theta trough, while mid-frequency gamma oscillations occur close to the theta peak (Supplementary information Fig S2C, D). This is opposite to the relationship reported by Colgin et al., 2009, but is similar to what is reported by other work in rats [14] and mouse [16, 27]. While input-specific oscillatory components in CA1 can be de-mixed using high-density silicon probe recordings with current source density (CSD) analysis [27] (see also Supplementary information Fig S2B, C) or independent component analysis [28], here we exploit that both slow and mid-frequency gamma oscillations can be identified in CA1 *stratum pyramidale*, which is both the target of place cell recordings, and the basis of virtually all the data upon which the routing-by-synchrony hypotheses are based.

We began by comparing power in representative frequency bands for slow gamma (30-50 Hz) and mid-frequency gamma (70-90 Hz) and their respective power ratio (Fig 1D). Before the mouse initiated successful avoidance movements, the ratio of slow to mid-frequency gamma power progressively increased from ˜5 s prior to the initiation of avoidance movements with the maximum ratio occurring ˜1 s prior to the active avoidance. This relationship was confirmed with one-way ANOVA (F_2,5168_ = 294.84; p = 5.6×10^-122^) of the differences between three time intervals (-7..-5 s, -2..0 s and 0..2 s) around the avoidance onset. Post-hoc Dunnett’s tests confirmed significant differences from the -7..-5 s baseline interval for intervals just before (-2..0 s) and just after (0..2 s) the initiation of active avoidance (p < 0.001 in both cases). The power ratio was strongly negatively correlated with speed (Fig 1D, bottom; Pearson’s correlation r = - 0.25, p = 0) as has been reported [29]. Because changes in speed confound associating these changes in the LFP with recollection, we examined alternative approaches for characterizing gamma changes in the LFP that are minimally impacted by speed and instead emphasize the internal cognitive information processing upon which the routing-by-synchrony hypothesis is based.

The routing-by-synchrony hypothesis also predicts that information between two networks is relayed most effectively during high-power, synchronized oscillatory states in contrast to all non-oscillatory activity which gives rise to the 1/f power spectra of LFP and EEG signals [30, see also Supplementary material Fig. S1B]. Because the present work relies on comparing oscillations of different frequency bands, to avoid potentially misleading estimates of the relative strength of oscillations from 1/f organized power spectra, we build on our prior work and discretized continuous LFP signals into frequency-specific oscillatory events and their rates [31]. Oscillatory events were detected as local power maxima in the z-score normalized, wavelet-transformed LFP signal (Supplemental information Fig S2E; also refer to Fig S3 for discussion about threshold setting for event detection). To compute rates of oscillatory events, we first selected representative frequency bands for slow gamma (30-50 Hz) and mid-frequency gamma events (70-90 Hz; refer to Supplemental information Fig S3 for discussion about band selection) and then computed event rates as the number of detected events in a given frequency range above 2.5 S.D. power at *stratum pyramidale* in 1000-ms long windows advanced by 250 ms consistent with prior routing-by-synchrony studies [7, 12, 32]. From now on, we therefore use slow (30-50 Hz) and mid-frequency (70-90 Hz) gamma event rates. These are defined as the number of detected oscillatory events in a 1-s long interval that averages several theta cycles, and avoids potential controversies around which oscillations dominate a single theta cycle. We compute their respective ratios, SG/MG as the ratio of the slow to mid-frequency gamma event rates and MG/SG as the ratio of mid-frequency to slow gamma event rates.

### The ratio of slow to mid-frequency gamma oscillation events is maximal when recollection likelihood is high

The session-specific slow gamma (SG) and mid-frequency gamma (MG) oscillation rates and the SG/MG ratio were examined around the time of successful avoidances of the initial location of shock. The mid-frequency gamma oscillation rate decreased with the minimum occurring 2 – 0.5 s before avoidance onset (Fig 1F, left; compare to wavelet spectrum in Fig 1E). Slow gamma had a less pronounced decrease and could even increase before avoidance onset. Slow gamma increased after avoidance onset, peaking about 500 ms afterwards, preceding mid-frequency gamma, which peaked at 750 ms. In contrast to the power ratio (Fig 1D), the SG/MG ratio was only weakly correlated with speed (SG/MG ratio: r = - 0.09, p = 4.6×10^-22^, explaining < 1% of the variance; power ratio: r = -0.25, p = 0, explaining > 6% of the variance; t test for difference between means of Fisher-transformed correlations t_16_ = 2.13, p = 0.048). The SG/MG ratio was maximal 1-2 seconds before and it was minimal ˜1 s after avoidance onset (Fig 1F, left). These relationships were confirmed with one-way ANOVA (F_2,4134_ = 54.22; p = 5.7×10^-24^) on the SG/MG ratios between three time intervals (-5..-3 s, -2..0 s and 0..2 s) around avoidance onset. Post-hoc Dunnett’s test confirmed significant differences from the -5..-3 s baseline interval, for intervals just before (-2..0 s) and just after (0..2 s) the initiation of active avoidance (p < 0.001 in both cases). The comparison of the two intervals before entering the shock zone was not significant (F_1,79_ = 0.38; p = 0.54) when the mouse failed to actively avoid shock but nonetheless initiated running away from the shock zone upon being shocked (Fig 1F, right). Because this slow gamma dominance over mid-frequency gamma (SG_dom_) was identified during a few seconds of stillness prior to avoidance of the shock location, it is possible that SG_dom_ either indicates momentary recollection of the shock locations, or preparation (initiation) of locomotion.

### Slow gamma dominance predicts successful place avoidance

We then investigated whether the rates of slow or mid-frequency gamma oscillations, or their ratio indexed behavior potentially associated with recollection or initiation of movement *per se.* We first examined the time series of the SG/MG ratio but did not restrict the analysis to peri-avoidance episodes with preceding stillness (Fig 2). To compute time intervals between the SG/MG maxima that define SG_dom_, we first detected local peaks in the SG/MG ratio series with amplitude > 1 (i.e. SG > MG) and then selected the subset of maxima with prominence (amplitude difference between maxima to the preceding and following minima) > 1. This step excluded short intervals resulting in multiple peaks in a sequence (see Fig 2A).

**Fig 2.**
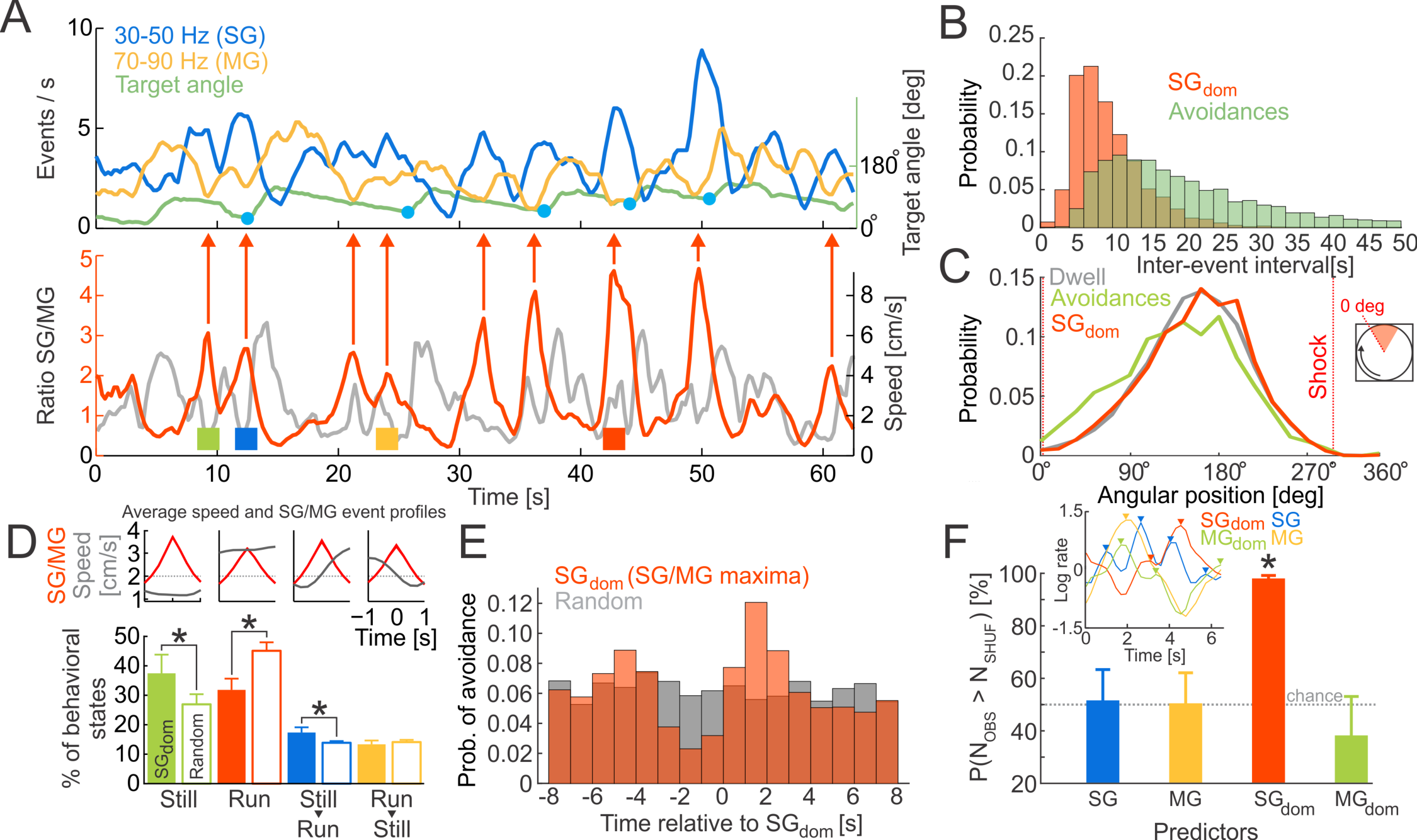
Slow gamma dominance predicts active avoidance. (A) Top: time series of slow gamma and mid-frequency gamma event rates and the angular distance of a mouse from the leading edge of the shock zone. Avoidances are marked by blue dots. The leading edge of the shock zone corresponds to 0°. Bottom: time series of slow to mid-frequency gamma ratio (SG/MG) with local maxima (SG_dom_) indicated (red arrows). (B) Probability distributions of inter-event intervals for consecutive SG_dom_ events and successful avoidances during training sessions. (C) Angular distributions of a mouse’s location (Dwell; gray), locations of avoidances (Avoidances; green), and locations of SG_dom_ events (SG_dom_; red). (D) Proportions of different behavioral events detected during SG_dom_ events (filled bars) and randomly selected events (empty bars). Average SG/MG ratio and speed in 2-s windows centered around SG/FG maxima is shown at the top. Dotted line represents the speed threshold 2 cm/s used for the behavior classification. Corresponding examples of behavioral states are marked by colored squares in A, bottom. *p < 0.01. (E) Probability of observing an avoidance relative to a SG_dom_ event and randomly selected times. (F) The probability of predicting avoidances by chance (after randomly shifting the time stamps of detected maxima), by using the maxima of the slow gamma, the mid-frequency gamma, SG_dom_ events or the MG/SG ratio maxima (MG_dom_ events). The inset shows examples of detected maxima in the four series types. *p < 0.05 relative to chance. Data are represented as average ± SEM.

We first investigated whether the occurrence of SG_dom_ events (avg. inter-event time 9.3 s) and avoidances (avg. inter-event time 26.0 s; Fig 2B) were substantially similar or different (compare upper and lower time series in Fig 2A). SG_dom_ events were more frequent than avoidances (Kolmogorov-Smirnov test D_3433_ = 0.522, p = 3.2×10^-8^) indicating that not every SG_dom_ event is followed by avoidance behavior. This means that while SG dominance in the CA1 LFP was initially identified by focusing on peri-avoidance episodes defined by stillness changing to locomotion, a parsimonious account for SG dominance is it is more likely to indicate moments of active recollection than just preparation for initiating movement (see Supplementary information Video S1).

We next analyzed the length of SG_dom_ events by thresholding the SG/MG ratio time series. The SG_dom_ peaks lasted 3.0 ± 2.7 s when the threshold was SG/MG ≥ 1 and they lasted 1.2 ± 1.1 s, when the threshold was SG/MG ≥ 2, suggesting that SG dominance likely lasts several theta cycles.

We then assessed the spatial distribution of SG_dom_ events (Fig 2C). Consistent with these being internal, cognitive events, the spatial distribution of the SG_dom_ events resembles the spatial distribution of where the mice visited (maximal dwell opposite the shock zone; Mann-Whitney-Wilcoxon non-parametric test compared to dwell U = 2.8×10^9^, p = 0.43) and accordingly, these places differed from the places where the mice express avoidance behavior by initiating movement away from the leading edge of the shock zone (Mann-Whitney-Wilcoxon non-parametric test compared to dwell U = 9.4×10^8^, p = 0). These data are consistent with SG_dom_ being related to an internal cognitive variable like active recollection, where recollection might not only be of the locations of shock.

We next studied in what behavioral states SG_dom_ events occur (Fig 2D). We classified behavioral states using each mouse’s average speed during -1..-0.25 s before and 0.25..1 s after SG_dom_ events. The “Run” state had average speed before and after a SG_dom_ event > 2 cm/s. The “Still” state had average speed before and after a SG_dom_ event < 2 cm/s. “Still->Run” state had average speed before a SG_dom_ event < 2 cm/s and after the SG_dom_ event ≥ 2 cm/s. “Run->Still” state had average speed before a SG_dom_ event ≥ 2 cm/s and after the SG_dom_ event < 2 cm/s.

SG_dom_ events occur during both active movement and stillness. During pretraining recordings when the mice explored the rotating arena prior to ever experiencing shock, the majority (˜75%) of observations during SG_dom_ or random events comprised continuous stillness or running with greater prevalence of running. The prevalence of these movement-defined states were indistinguishable during SG dominance and randomly selected episodes (Supplementary information Fig S4B). Place avoidance training changed which movement-defined behaviors are expressed during SG dominance. Overall, the continuous stillness and running behaviors still account for about 70% of observations during SG_dom_ events; transitional behaviors from stillness to running or *vice versa* are less frequent (Fig 2D; F_3,35_ = 8.88, p = 0.0002; post-hoc Dunnett’s test against Still: Still = Run > Still→Run > Run→Still). We then computed the frequencies of observing these movement-defined behaviors during the same number of random intervals as were identified for SG_dom_ (empty bars in Fig 2D). Overall, the majority (˜75%) of observations were during continuous stillness or running like during SG dominance. However, the SG_dom_ and random event comparisons indicate that Stillness and transitions from Stillness to Running are overrepresented during SG_dom_, while Running is underrepresented during SG_dom_ events (χ^2^ test for multiple proportions χ^2^_3_ =119.1, p = 8.4×10^-25^; Still: χ^2^_1_ = 36.4, p = 2.4×10^-7^; Run: χ^2^_1_ = 62.6, p = 8.1×10^-13^; Still→Run: χ^2^_1_ = 19.9, p = 0.0005; Run→Still: *χ*^2^_1_ = 0.15, p = 0.99). Thus, prior to the place learning task, the prevalence of movement-related behaviors is indistinguishable from chance during SG dominance, but the prevalence of these behaviors deviates from chance to favor behaviors that are associated with high likelihood of recollecting the location in which a shock was previously experienced. These investigations of the prevalence of SG_dom_ during movement-defined behaviors indicate it is unlikely that SG_dom_ can be fully explained by movement planning or initiation.

To further evaluate the possibility that SG dominance is indicative of long-term memory recollection, we tested the ability of the SG_dom_ events to predict successful avoidances, reasoning that recollecting locations of shock should precede effective avoidance behavior. First, we examined the probability of observing an avoidance at times relative to SG_dom_ events and compared that distribution to the probability of observing an avoidance at times relative to randomly-selected events. The distributions were different and there was an increased occurrence of avoidances 1-2 seconds after SG_dom_ events (Fig 2E; Kolmogorov-Smirnov test D_2715_ = 0.08, p = 1.6×10^-8^). Second, we created four avoidance predictors that used either the maxima in SG rate, maxima in MG rate, maxima in SG/MG ratio (i.e. SG_dom_) or maxima in the MG/SG ratio (i.e. MG_dom_). Prior to detecting these peaks, the ratios (SG/MG and MG/SG) were log-transformed and all time series were z-score normalized and only maxima with z-score values > 0.5 S.D were selected to guarantee similar rates of detected peaks in all four time series. Avoidances were predicted in a 4-s long window following the maxima. Note that even though the MG/SG ratio is the inverse of the SG/MG ratio, the maxima (i.e. SG_dom_ and MG_dom_) in both series occur at different times (Fig 2F inset). The four maxima types differed in their ability to predict avoidance (Fig 2F; F_3,43_ = 10.5, p = 2.0×10^-4^); only SG_dom_ had predictive power better than chance (t_8_ = 24.56; p = 4.0×10^9^). While SG dominance occurred regularly and everywhere and during active and passive behavioral states, it nonetheless predicts successful place avoidance in the immediate future, consistent with SG_dom_ signaling recollection of long-term memories.

### Abnormal recollection in Fmr1-KO mice predicts excessive slow gamma dominance

We investigated the hypothesis that SG_dom_ events indicate long-term memory recollection by taking advantage of prior work with Fmr1-knockout (KO) mice [33]. These mice express a null form of the Fmr1 gene to model the genetic defect in FXS, a syndromic form of autism and the most common inherited form of intellectual disability [34]. Place avoidance learning and 24-h retention of long-term memory for the initial shock zone location appears normal in Fmr1-KO mice, but Fmr1-KO mice express cognitive inflexibility when they must avoid the formerly preferred place because the shock is relocated 180° on a conflict test [33]. We replicated this observation in the mice we recorded (Fig 3A; time spent in six 60° wide spatial bins during the second half of the conflict session: Genotype: F_1,14_ = 4.41, p = 0.05; Bin: F_5,10_ = 64.26, p = 2.7×10^-7^; Genotype x Bin: F_5,10_ = 2.65, p = 0.09). Whereas wild-type mice quickly adapt to the new location of shock on the conflict session, Fmr1-KO mice are impaired, possibly because they persist in recalling the former shock location that is now incorrect (Fig 3B; Genotype: F_1,44_ = 6.96, p = 0.01; Session: F_1,44_ = 77.32, p = 3.0×10^-11^; Time: F_1,44_ = 48.62, p = 1.2×10^-8^; Genotype x Session x Time: F_1,44_ = 11.16, p = 0.002; Post-hoc tests confirm that WT and Fmr1-KO only differ in the second half of the conflict session).

**Fig 3.**
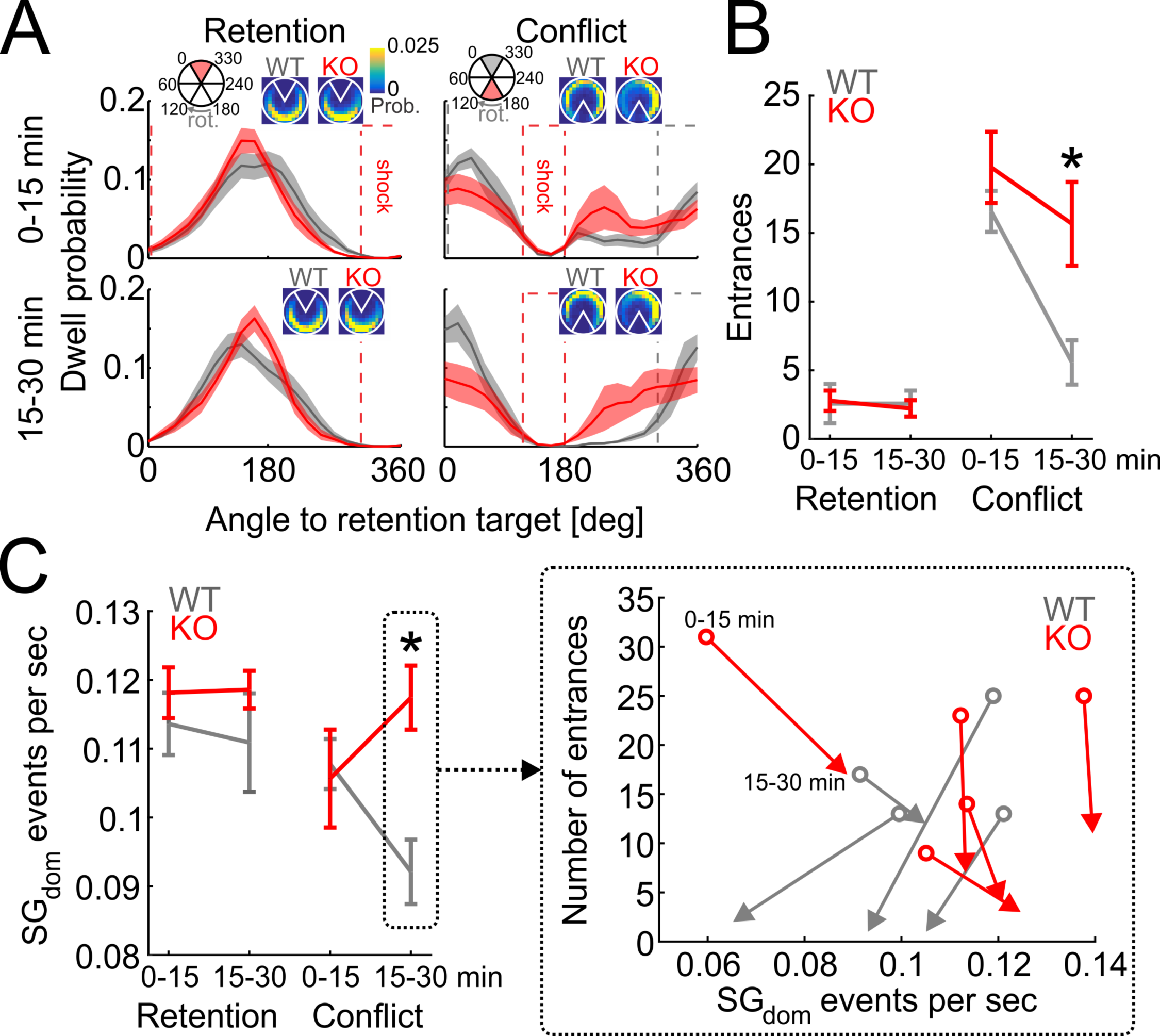
Cognitive inflexibility and associated increases of slow gamma dominance in Fmr1 KO mice. (A) Dwell distribution during first half (0-15 min; top) and second half (15-30 min; bottom) of retention (left) and conflict (right) sessions for wild-type (gray) and Fmr1-KO (red) mice. Dotted lines show locations of the active shock zone during each session (red) and location of the initial shock zone during conflict sessions (gray). Insets show dwell probability distributions. (B) Behavioral performance during retention and conflict sessions for wild-type and Fmr1-KO mice. (C) Rates of SG_dom_ events during retention and conflict sessions. *p < 0.05 between genotypes. Data represent average ± SEM. Inset: vectors showing the time-evolution from the first half (circles, 0-15 min) to the second half (arrowheads, 15-30 min) of the conflict session in the coordinate system of x = SG_dom_ rate and y = number of entrances.

We then examined if SG dominance distinguishes the wild-type and Fmr1-KO mice in the conflict session when the mutants express inflexibility compared to the wild-type mice. During the initial half of the conflict session when the genotypes are behaviorally similar, the rate of SG_dom_ events was also indistinguishable between the genotypes. However, the wild-type SG_dom_ rate decreased in the second half of the conflict session while the Fmr1-KO rate increased, resulting in a significant Genotype x Time interaction (F_1,24_ = 5.59, p = 0.027; Fig 3C), and a marginal Genotype x Session x Time interaction (F_1,24_ = 3.52, p = 0.07) because the genotypes only differed on the second half of the conflict session when wild-type mice decreased both the number of errors and the rate of SG_dom_ events but the mutant mice did not (Fig. 3C right detail), no other effects were significant (F_1,24_′s ≤ 1.71, p′s > 0.2). These findings are consistent with the idea that SG dominance reflects recollection of long-term memories, and suggest the possibility that Fmr1-KO mice may express abnormally persistent recollection of conditioned place avoidance memory.

### Slow gamma dominance predicts non-local place coding in neural discharge during active place avoidance

Next, based on evidence that place cell discharge is more likely to represent non-local, distant places during recollection [35-37], we tested the hypothesis that SG dominance identifies recollection. The hypothesis predicts that during the place avoidance task, putative pyramidal cell discharge is non-local during SG dominance, assessed as increased error in the location estimate obtained from ensemble firing rates using a Bayesian decoder [38]. We examined CA1 putative pyramidal cell discharge from four wild-type and three Fmr1-KO mice after initial and conflict avoidance training. For these analyses, the SG_dom_ events were detected independently from all LFP signals recorded at tetrodes on which putative pyramidal cells were identified. We made this decision to avoid bias by choosing one of the tetrodes as representative because detection of SG_dom_ events on all pairs of tetrodes in a given animal was coincident 25% (24.1±18.4% in wild-type and 25.1±15.8% in Fmr1-KO mice). As predicted, during SG dominance, CA1 putative pyramidal cell discharge decodes to distant locations (Fig 4). The putative pyramidal cell ensemble recorded during active place avoidance (Fig 4A) shows in example Fig 4B, that the error between the observed and estimated locations is increased during the SG_dom_ events just prior to avoidance behavior. We computed the average decoding error time-locked to the SG_dom_ events, during which we hypothesize recollection. The decoded error is the z-score normalized average of the 1-D posterior probability multiplied by the error function, which was 0 at the observed 1-D angular location and linearly increased with the distance from the observed location. For comparison, the decoding error was also computed time-locked to random moments as well as relative to MG_dom_ events (Fig 4C). The average decoding error was indistinguishable between the genotypes (wild-type: 47.5 ± 3.6 degrees; Fmr1-KO: 52.0 ± 4.8 degrees; t_10_ = 0.77, p = 0.46). The Bayesian decoding error in both wild-type and Fmr1-KO mice was large around the time of SG_dom_ events, in contrast to the relatively small error associated with random times and with MG_dom_ events, during which the error was minimal (Fig 4D). Indeed, the decoding errors were greatest during SG_dom_ events (F_2,4215_ = 7.87, p = 4×10^-4^; Dunnett’s test: SG_dom_ > MG_dom_ = RND), and although this pattern appeared more extreme in Fmr1-KO mice at the time of the event, place representations in Fmr1-KO ensemble discharge did not differ from wild type (Genotype: F_1,4215_ = 0.04, p = 0.9; Genotype X Event Interaction: F_2,4215_ = 0.33, p = 0.72). This result could arise because the Bayesian posterior during SG_dom_ events is less localized and thus more imprecise, or alternatively, during SG_dom_ events the posterior could be just as compact as during non-SG_dom_ moments when putative pyramidal cell discharge decodes to current locations.

**Fig 4.**
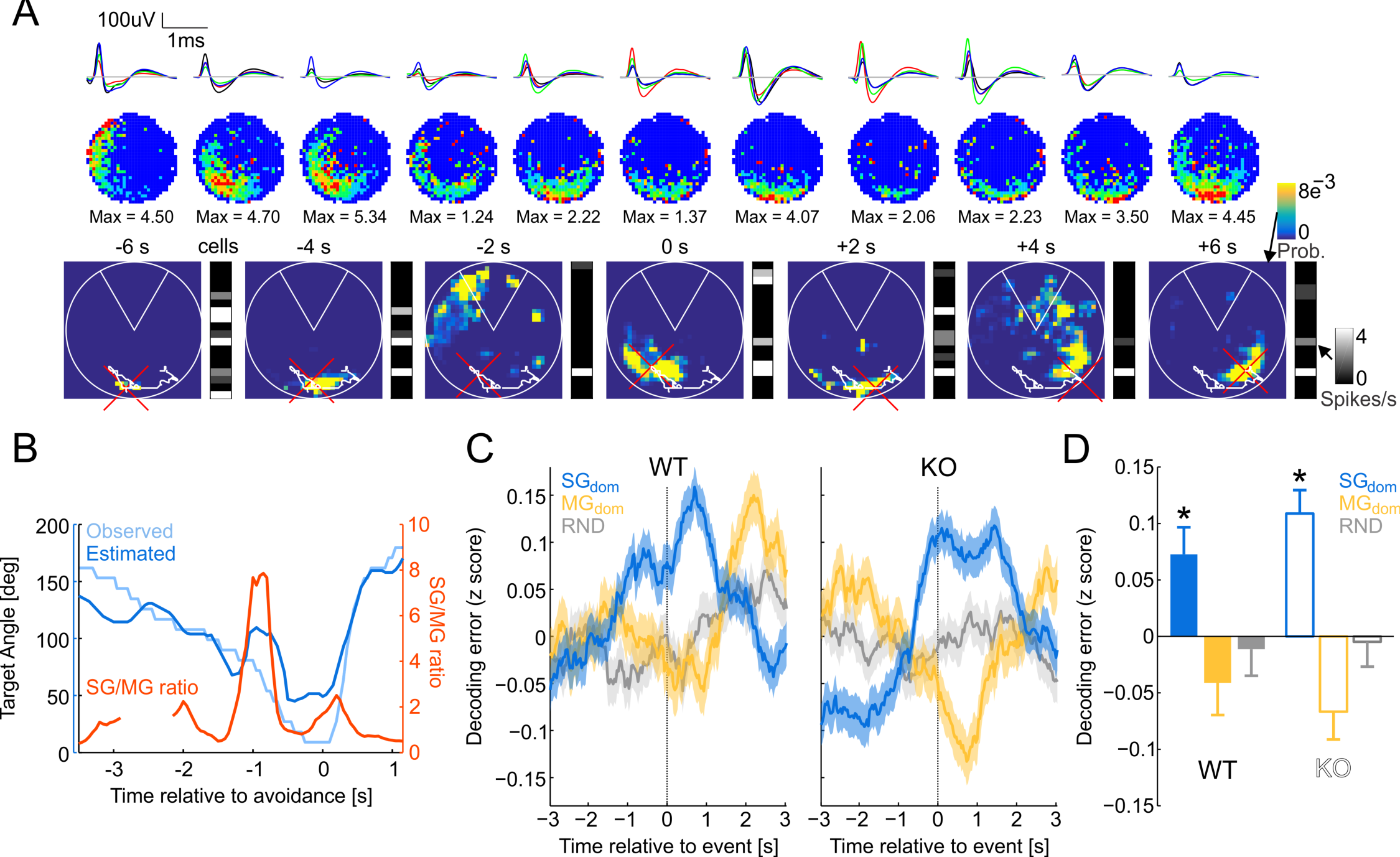
Error in Bayesian decoding of location increases during SG_dom_ events. (A) Example firing rate maps (top) and 2-D decoded Bayesian posterior around avoidance onset. Ensemble activity vectors are shown to the right of each decoded Bayesian posterior. The mouse’s path during a 12-s segment is shown as a white line and the current position is marked by a red cross. (B) Example time series of the angular position that was observed and decoded using a 1-D Bayesian estimator from ensemble discharge overlaid with the SG/MG ratio. Time T = 0 s marks avoidance onset. (C) The average of wildtype (WT) and Fmr1 KO (KO) z-score normalized 1-D decoding error from ensemble activity that is time-locked to SG_dom_ events, MG_dom_ events, and random times (RND). Time T = 0 s corresponds to the time of the events. (D) Summary of decoding error at the moments of SG_dom_ events, MG_dom_ events and random times for WT and KO mice. *p < 0.05 relative to random. Data are represented as average ± SEM.

To control for the possibility that the mouse’s speed might differ during SG_dom_ and random events, we restricted the analysis only to times of stillness (speed < 2 cm/s; 33% of SG_dom_ events) the same pattern was observed although the effect of genotype became significant because the decoding error was higher in WT during SG_dom_ events, while decoding error was lower in Fmr1-KO during MG_dom_ events (genotype x oscillation type 2-way ANOVA, genotype: F_1,1387_ = 8.38, p = 0.004, oscillation type: F_2,1387_ = 22.23, p = 3.1×10^-10^, interaction: F_2,1387_ = 0.03, p = 0.98; Dunnett’s test for difference from random events: SG_dom_: p = 0.008, MG_dom_: p = 0.049). When analysis was restricted to the times during running (speed ≥ 2 cm/s; 56% of SG_dom_ events), we observed only genetic differences, because Fmr1-KO mice show both higher error of decoding during both SG_dom_ and MG_dom_ events, while wild-type mice show the same pattern of increased decoding error during SG_dom_ and reduced decoding error during MG_dom_ events (genotype x oscillation type 2-way ANOVA, genotype: F_1,2353_ = 7.20, p = 0.0073, oscillation type: F_2,2353_ = 0.81, p = 0.45, interaction: F_2,2353_ = 0.39, p = 0.68). The size of the posteriors were indistinguishable during SG_dom_, MG_dom_, and random moments when we decoded 2-D position (F_2,2127_ = 0.74, p = 0.47). In fact, the posteriors were most compact during SG_dom_ when we decoded the mouse’s 1-D angle in the arena relative to the leading edge of the shock zone (F_2,2127_ = 5.04, p = 0.006; SG_dom_ < MG_dom_ = RND according to post hoc Dunnett’s tests), indicating the nonlocal representations of position during SG_dom_ were compact and precise. These findings confirm that pyramidal cell ensemble discharge selectively represents distant locations during SG dominance, consistent with recollection of locations remote from the mouse.

Because of the role of sharp-wave ripple (SWR) events in the replay of non-local place cell sequences [39], including during fear memory expression [40], we further investigated this non-local decoding during isolated SG events (events detected in the 30-50 Hz band without concurrent MG or SWR events). For comparison, we also investigated isolated MG events (events detected in the 70-90 Hz band without concurrent SG or SWR events; see Supplementary information Fig S5). This excluded the approximately 10% of SG and MG events that were concurrent with SWR events in both wild-type and Fmr1-KO mice (Supplemental information Fig S5A). Both wild-type and Fmr1-KO putative pyramidal cell representations appeared more non-local during SG events compared to MG events, indicating that events during sharp wave ripples cannot account for the observation that SG_dom_ is associated with nonlocal hippocampus place representations (Supplementary information Fig S5C, D; Genotype: F_1,117515_ = 22.57, p = 2.0×10^-6^; Oscillation: F_2,117515_ = 34.40, p = 1.2×10^-15^; Genotype x Oscillation Interaction: F_2,117515_ = 3.27, p = 0.037; Dunnett’s test for difference from random events: SG_dom_: p < 0.0001, MG_dom_: p < 0.0001). These statistical tests included the ensemble firing rate as a covariate because of the significant relationships between decoding error and putative pyramidal cell firing rates (WT : r^2^ = 9%, p = 0; Fmr1-KO: r^2^ = 13%, p = 0), whereas the relationships to speed explained substantially less of the variance (WT : r^2^ = 0.0003%, p = 0.69; Fmr1-KO: r^2^ = 0.4%, p = 2.7×10^-35^). These findings with isolated slow gamma events, as well as those with slow gamma dominance, suggest that place memory recollection is predicted by slow gamma dominance, which also identifies when pyramidal cell ensembles will represent remote places, consistent with the hypothesis that slow gamma dominance in hippocampus CA1 identifies active recollection of long-term memory.

### Putative pyramidal cell discharge during slow gamma dominance represents places that will be avoided

Finally, we analyzed the Bayesian posterior probability maps from the decoding to examine whether during avoidance sessions, putative pyramidal cell representations during SG dominance decode to the vicinity of the shock that the mouse will avoid, consistent with recollection of the places to avoid. Fig 5A shows four example Bayesian 2-D posterior probability maps computed at times before to times after individual avoidances. There are two examples from each genotype, one when the shock was in the initial location and the other after a conflict session with relocated shock. These examples illustrate that up until ˜2 s before the avoidance, the peak values of the posterior probability correspond to the mouse’s location. However, ˜2 s before the avoidance of the initial shock location, in both genotypes, the posterior probability can peak at non-local positions that are in the vicinity of the shock zone or 180° away, which is the safest and most frequented location on the arena during training to the initial shock location. The genotypes differ in the conflict session, remarkably. The wild-type example shows non-local decoding to the currently-correct, relocated shock location ˜2 s before avoidance, whereas the Fmr1-KO example shows decoding to the currently-incorrect shock location that was formerly correct.

**Fig 5.**
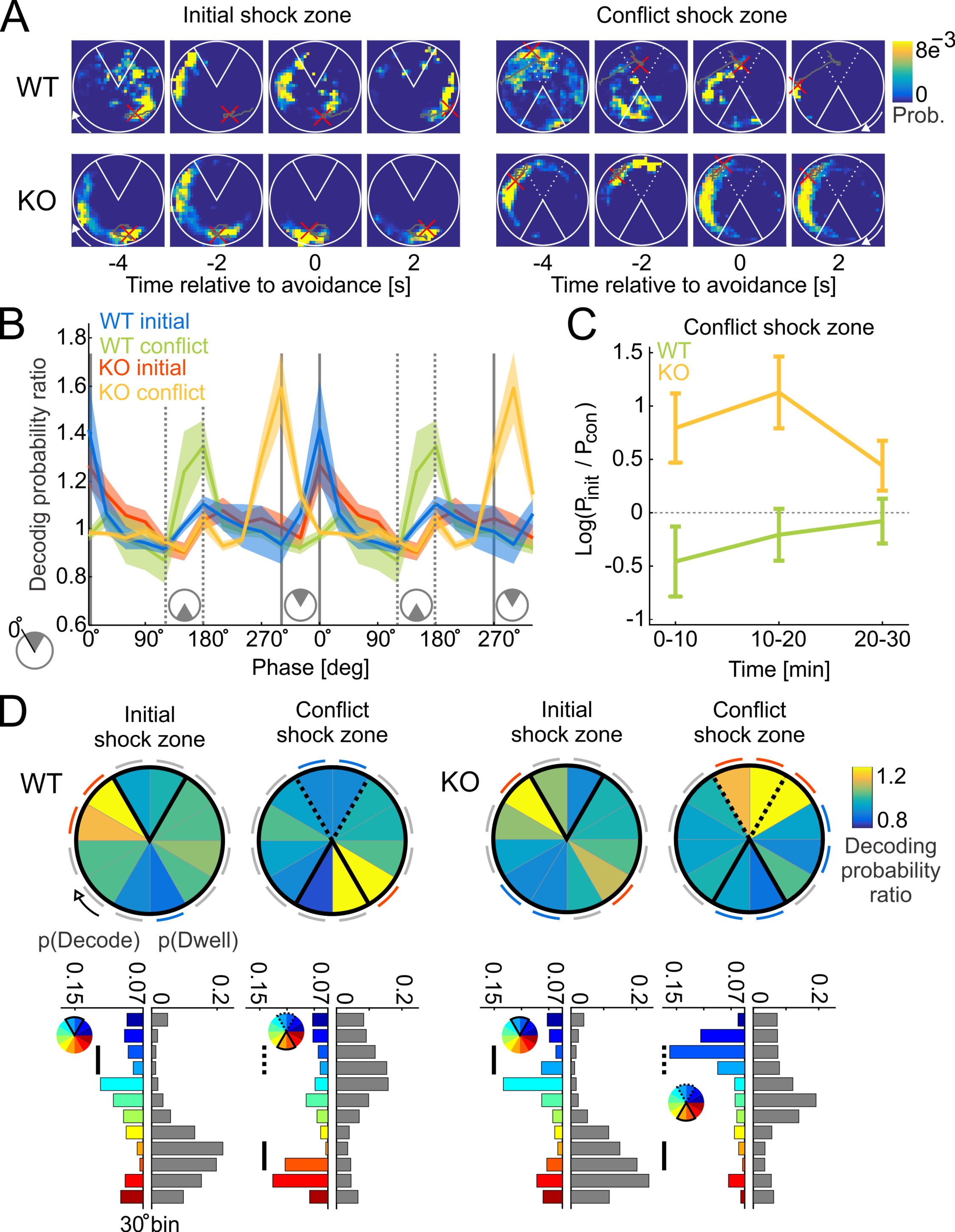
During SG_dom_ events, putative pyramidal cell ensemble discharge represent the vicinity of shock. (A) Four examples of the time evolution of the Bayesian posterior probability from before to after avoiding the shock zone (white sector centered at 12 o’clock for the initial shock zone and 6 o’clock for the conflict shock zone). The mouse’s path during the episode is shown in gray with the current location indicated by a red cross. Top row corresponds to a wild-type mouse, bottom row to a Fmr1-KO mouse. The left examples illustrate training to the initial shock zone. The right examples are after the shock was relocated for conflict training. The initial shock zone location in conflict training is shown as a dotted line. (B) Average normalized posterior probability as a distance from the leading edge of the shock zone. Full gray lines mark the location of the initial shock zone, dotted lines mark location of the conflict shock zone. Data are represented as averages ± SEM. (C) Ratio of average posterior probability at the location of the initial shock zone and the location of conflict shock zone in 10-min intervals. (D) Top: summary of normalized posterior probability estimates obtained during SG_dom_ events for the initial and conflict shock zone sessions for wild-type (left) and Fmr1-KO (right) mice. Notice maximal decoding probability at the leading edge of the shock zone in both WT and KO mice during the initial shock zone session, and during the conflict session, maximal decoding probability is at the leading edge of the relocated shock zone in WT mice but not in Fmr1-KO mice. During conflict, Fmr1-KO ensemble discharge decodes to the vicinity of the initial shock zone that is currently incorrect for avoiding shock. Red arcs located next to angular bins indicate significantly positive (>1) normalized probability (p < 0.05), blue arcs indicate significantly negative (<1) normalized probability (p < 0.05), gray arcs indicate n.s. relative to 1. Bottom: Split bar plots comparing the decoding probability distributions (color; each color corresponds to one 30° angular position) and dwell distributions (gray) for corresponding trials. Black vertical lines mark regions inside of the currently correct shock zone. Dotted vertical lines in conflict trials mark regions inside the initially correct shock zone.

Similar patterns of representational flexibility in wild-type and inflexibility in Fmr1-KO are seen in the summary data, computed as the ratio of the posterior during SG_dom_ events normalized by the average posterior during MG_dom_ events, when decoding was local. During SG dominance, this posterior ratio peaks in the vicinity of the initial location of shock and this is observed for both wild-type and Fmr1-KO putative pyramidal cell representations (blue and red data in Fig 5B, respectively). The posterior ratio peaks in the vicinity of the currently-correct location of shock in the post-conflict session, but only for wild-type putative pyramidal cell representations, demonstrating representational flexibility (green data in Fig 5B). In the post-conflict session, the Fmr1-KO posterior ratio peaks adjacent to the currently-incorrect shock zone (yellow data in Fig 5B). The SG_dom_/MG_dom_ posterior probability ratio averaged across the location of the initial shock zone and the location of the conflict shock zone shows higher decoding probability of the initial shock zone for Fmr1-KO and higher decoding probability towards the conflict shock zone in the wild-type ensembles (Fig 5C; Two-way ANOVA, genotype: F_1,1000_ = 15.44, p = 9.1×10^-5^; interval: F_2,1000_ = 0.56, p = 0.57; genotype x interval: F_2,1000_ = 1.17, p = 0.31). The difference in decoding probability became similar in the two genotypes during the 20-30 min interval, consistent with the mice ceasing avoidance behavior when the avoidance is not reinforced. Statistical comparisons of the putative pyramidal cell posterior ratios confirm significant overrepresentation of the regions adjacent to the leading edge of the initial shock zone in the wild-type and Fmr1-KO mice. Overrepresentation is also observed at the relocated shock location in the conflict sessions, but only in wild-type putative pyramidal cell representations (Fig 5D). Whereas, in Fmr1-KO putative pyramidal cell ensemble representations, the posterior ratios are overrepresented in the currently-incorrect shock location during the post-conflict sessions, as confirmed by the significant Genotype x Trial x Region interaction: F_11,36983_ = 4.15, p = 3.7×10^-6^ (the main effects of Genotype and Trial were not significant but the effect of Region was significant F_11,36983_ = 3.68, p = 3.0×10^-5^, the Genotype x Region interaction F_11,36983_ = 2.59, p = 0.0027 and Trial x Region interactions F_11,36983_ = 4.47, p = 2.98×10^-5^ were significant as well). Because avoidance behavior of Fmr1-KO and wild-type mice differed during the conflict sessions (Fig 3A), we also tested the possibility that overrepresentation of the currently incorrect shock zone in Fmr1-KO might be caused by corresponding overrepresentations of where the mice visited. We compared the angular probability distributions of the SG_dom_/MG_dom_ decoding probability ratios and the dwell distributions in the sessions with the initial and conflict shock zones (Fig 5D, bottom), by computing a t-distributed test statistic as the absolute difference between the decoding and dwell distributions divided by the average absolute difference between the first and second halves of each of the decoding and dwell distributions; initial shock zone: WT: t_11_=3.54, p=0.002; Fmr1-KO: t_11_=2.20, p=0.02; conflict shock zone: WT: t_11_=1.80, p=0.04; Fmr1-KO: t_11_=2.02, p=0.03). These findings demonstrate that SG dominance corresponds to activation of non-local, memory-related pyramidal cell representations and demonstrate for the first time, representational inflexibility in Fmr1-KO mice concurrent with behavioral inflexibility (Fig 3).

## DISCUSSION

### Summary – a neurophysiological hypothesis for recollection

To identify a neural signature of recollection, we began by selecting an enriched sample of potential recollection events using behavioral criteria (Fig 1) and investigated the rate of occurrence of gamma oscillations in the dorsal CA1 LFP. Individual slow and mid-frequency gamma events were not predictive, but SG dominance, i.e. maxima in the slow/mid-frequency ratio of event rates at *stratum pyramidale* predicted successful place avoidance (Fig 2), suggesting SG dominance is a candidate neural signature of long-term memory recollection, at least for place memories. While wild-type mice attenuated SG dominance when it was necessary to suppress recollection of the initially learned locations of shock, SG dominance was not attenuated in Fmr1-KO mice when they demonstrate cognitive inflexibility (Fig 3). SG dominance occurred approximately every 9 s in standard conditions of exploration as well as during place avoidance sessions, which is almost three times more frequent than active avoidance-like behaviors (Fig 2). This indicates that if SG dominance corresponds to an internal variable like recollection, then it may not merely be the recollection of conditioning events such as locations of shock. Indeed, SG dominance coincides with non-local putative pyramidal cell representations during post-avoidance recording sessions (Fig 4). We note that although Fmr1-KO mice model the genetic defect in FXS and express a number of biochemical and synaptic abnormalities [41, 42] their putative pyramidal cells that are classified as place cells express normal place fields [43], which makes their cognitive flexibility deficits a challenge to explain. However, guided by the notion that SG dominance identifies long-term place memory recollection, we observed neural representational inflexibility in Fmr1-KO mice, when they express behavioral inflexibility (Fig 5). Together, these findings provide convergent evidence that SG dominance predicts recollection as well as abnormalities of recollection in Fmr1-KO mice.

### Summary – Considering Alternative Interpretations

Recollection is an internal variable, inaccessible to direct observation and so alternative accounts for SG dominance were considered, but none are compatible with all the observations. The findings are incompatible with the alternative possibility that SG dominance merely indicates a process that anticipates or prepares to initiate movement (Fig 2). Movement preparation might have appeared to account for SG dominance because we initially identified SG_dom_ events by assessing avoidance behavior that was characterized by a period of stillness before the mouse moved away from the location of shock. However, after recognizing that SG dominance might be a neural correlate of recollection we investigated alternative accounts like avoidance behavior itself, movement preparation and other speed related possibilities. While successful avoidances are most frequent ˜1.5 s after SG_dom_ events (Fig 2E), SG_dom_ events are far more frequent and distributed distinctively from active avoidance movements (Fig 2B). Although SG_dom_ events are similarly present during continuous immobility and running, their likelihood is overrepresented during stillness and transitions from stillness to running compared to randomly selected moments (Fig 2D). During pretraining, before the mice experienced shock, the prevalence of movement-related behaviors during SG_dom_ events did not differ from the prevalence of these behaviors during random events (Supplemental information Fig S4B). Thus mere movement-related classifications of behavior do not account for when SG dominance is expressed. The deviations from chance expectations after place memory training, suggests that an internal cognitive variable may be influencing the expression of SG dominance. Nonetheless, because movement conditions vary with SG dominance, we considered other known movement related effects on gamma oscillations. Gamma frequency increases with running speed [44], more for faster than slow gamma frequencies [45], but SG_dom_ is distinctive; characterized by relatively increased rather than decreased SG power (Fig 1D). During steady immobility prior to avoidance movements, slow and fast gamma oscillations change and they change differently, indicating any relationship to speed is complex and not explained by known relationships to speed (Supplementary information Fig S3E). Similarly, because SG_dom_ events are associated with non-local place representations (Fig 4, Supplementary information Fig S5) they are unlike the events during the recently described N-waves that are associated with local place representations during immobility [46]. Furthermore, after isolating slow-gamma oscillatory events from contamination by mid-frequency gamma events and sharp-wave ripples and correcting for firing rate bias of the decoding, slow gamma events still expressed non-local decoding. Correspondingly, local place representations are predominantly observed during MG_dom_ events when SG is relatively weak (Fig 4; Supplementary information Fig S5). Consequently, SG_dom_ events may represent a complementary network state, in particular because, unlike N-waves, the discharge associated with SG_dom_ events is nonlocal (Fig 4, Supplementary information Fig S5), consistent with SG dominance signaling recollection of long-term memories of remote places and/or spatial events. By inspection, despite the non-local hippocampal representational discharge, vicarious trial-and-error [47-49] did not coincide with SG dominance. This was confirmed by statistical comparisons of numerical estimates of VTE [49] during SG dominance, MG dominance, and random events (Supplementary information Fig S6G; ANOVA F_2,9857_ = 0. 1, p = 0.9). Putative pyramidal cell discharge is also non-local during sharp-wave associated ripple events during which sequences of place cell discharge from active behavior can be observed to replay [50-53]. This sharp-wave associated replay is thought to underlie memory consolidation and support memory and decision making during the initial stages of learning [46, 54, 55]. Because SG_dom_ events are distinct from this sharp-wave associated discharge (Supplementary Information Fig. S5), the SG_dom_ events represent different phenomena. Thus the present findings of SG dominance and its associations to cognitive variables are distinct from known relationships of gamma to hippocampal network phenomena, and to overt behavioral variables. Accordingly, based on the present findings, we contend that SG dominance in the CA1 *stratum pyramidale* LFP is a neural signal that the hippocampus network is in the functional mode of recollecting long-term memories that are used to guide behavior, as in standard tests of long-term memory.

### Implications for the routing by synchrony hypothesis

Identifying SG dominance as a neural signal of recollection was inspired by prior work that proposed slow gamma oscillations measured at *stratum pyramidale* correspond to recollection event-associated activity from the CA3 region into *stratum radiatum*, whereas the neocortical inputs to the *stratum lacunosum-moleculare* are associated with higher frequency gamma and carry information about what is currently being experienced for encoding [7, 12, 13]. This important idea offers a solution for how multiple types of place information might be routed to the hippocampus to be used judiciously [8, 36, 56], for example to solve place avoidance tasks on a rotating arena during which distinct representations of the same physical places are activated [57, 58]. However, careful analysis of the extracellular currents along the dendritic compartments of dorsal CA1 has not supported this basic proposition [59, 60]. CA1 gamma-organized spiking is not simply entrained to the gamma-organization of the inputs from CA3 and entorhinal cortex, and the discrete oscillatory events at the *stratum radiatum* and *stratum lacunosum-moleculare* compartments have frequency components that overlap in the slow and mid-frequency gamma frequency bands [Fig. S2; 14, 16, 27, 33, 61]. Furthermore, these inputs are also relatively tonic, which has been both estimated [62] and observed during behavior [63, 64]. One factor that might add difficulty when interpreting differences in the literature is that most studies assume steady state cognitive conditions, which is not the case during either place avoidance or the foraging tasks that are often used, despite physical steady state conditions [57, 58, 65-67]. By selecting cognitively homogeneous samples, we find that recollection of hippocampus-dependent, actionable information is marked by the perisomatic region of CA1 being dominated by slow gamma over mid-frequency gamma oscillations, as if the two signals are in continuous competition (see Supplementary information Fig S2). These SG dominance “recollection” events appear to require a relatively large decrease in mid-frequency gamma in coordination with a lesser or no decrease in slow gamma, possibly depending on task conditions (Fig 1). Because mid-frequency gamma-associated entorhinal inputs facilitate CA1 spiking [14, 16], these observations point to a role for regulation of feedforward inhibition in the competition between temporoammonic and Schaffer collateral inputs to CA1 [16, 27, 68-70]. The present observations suggest that recollection of long-term memory is a transient change in the balance of the two signals, that is rapidly followed by a reversal of the dominance of midfrequency gamma by slow gamma. This result contrasts previous studies suggesting the appearance of one or the other type of gamma during processes of encoding and retrieval [7, 12, 13]. Contradictory observations that slow and mid-frequency gamma are commonly observed in the same theta cycle [Fig. S3; 14, 27] can be explained by the analyses that show the previously used thresholds of oscillation power selects for separate slow- and mid-frequency preferring theta cycles (Supplementary information Fig S3). While the present data do not support the details of the routing-by-synchrony hypothesis as first proposed [12, 13], the present findings support the gist, but without common feed-forward conceptions. Rather, this work has revealed dynamical operations within near continuous arrival of oscillation associated inputs along the somato-dendritic compartments of CA1 [63, 64, 69]. This input engages excitation, inhibition, and disinhibition, and is integrated locally in dendrites, such that the discharge of CA1 principal cells occurs as if embedded within a local neural activity infrastructure from which their spiking can emerge when the local infrastructure permits, by its transient adjustments to create distinctive information processing modes, like encoding and recollection [61, 63, 71-75]. These transient adjustments appear to emerge through a complex interplay between local neural dynamics and afferent activity [56, 60, 69, 76], and while the rules of engagement for this competition remain unknown, they are neither merely, nor predominantly feed-forward, in part because the gamma oscillations are locally generated and not dependent on gamma-paced inputs especially in the mid-frequency gamma and high gamma frequency ranges [16, 27].

### A neural signature of recollection

The recollection events we identified as perisomatic SG_dom_ events are brief, and they recur after several seconds, which may be a candidate mechanism for the seconds-long overdispersion dynamics in place representations that have been observed in single unit place cell ensemble studies during both place responding and foraging behaviors with no explicit cognitive demand [57, 58, 65, 66, 77]. Because the time scales differ and the SG_dom_ events span several theta cycles, they are unlikely to be the same cognitive information processing mechanism that governs the sub-second dynamics of single place cell spiking that can be observed as rodents traverse a cell’s place field and is interpreted as a form of encoding and recollection [7, 60, 78]. Rather the SG_dom_ events suggest that cognitive information processing is intimately tied to the coordinated regulation of inhibition at the perisomatic region, and perhaps elsewhere, under the control of the distinct, anatomically-segregated information-carrying afferents to CA1 [56, 61, 79], although the anatomical segregation of inputs may not be a requirement [69]. Resembling neural control of incompatible behaviors in leech [80], the present findings demonstrate in gamma a specific, dynamic coordination of excitation and inhibition that controls the cognitive information processing that permits effective spatial cognition, whereas its discoordination is associated with cognitive inflexibility [33, 56, 81]. Alterations in this coordination account for the cognitive effort of animal subjects both when they demonstrate adaptive cognitive information processing [14, 82] and when they exhibit inflexible cognition, as was observed in both the neural signals and the behavior of the wild-type and the Fmr1 KO mouse model of FXS and autism-related intellectual disability [33].

### Experimental Procedures

All methods comply with the Public Health Service Policy on Humane Care and Use of Laboratory Animals and were approved by the New York University and State University of New York, Downstate Medical Center Institutional Animal Care and Use Committees. Because detailed methods have been described [31, 33], only brief descriptions are provided.

### Subjects

A total of 21 wild-type mice with a mixed C57BL6/FVB background were used as well as 20 Fmr1-KO mice carrying the Fmr1^tm1Cgr^ allele on the same mixed C57BL6/FVB background. The mutant mice were obtained from Jackson Laboratories (Bar Harbor, ME) to establish local colonies. The mice were 3 - 6 months old. LFP recordings and behavior from 16 wild-type and 17 Fmr1-KO mice animals were studied in [33]. Of those mice, 9 wild-type mice were recorded during avoidance training with electrodes localized in *stratum pyramidale* and used for analyses in Fig 1 and Fig 2. Seven wild-type mice and 9 Fmr1 KO mice were used for behavioral analyses of 24-h retention of place avoidance and conflict training (Fig 3). Four wild-type mice and 5 Fmr1 KO mice with electrodes localized in *stratum pyramidale* were recorded during conflict training and used in electrophysiological analysis in Fig 3. Four wild-type mice and three Fmr1 KO mice were implanted with tetrodes and recorded after place avoidance training. These were used for the analyses in Figs. 4 and 5. One mouse was used for the CSD analysis in Supplementary information Fig S2.

### Surgery to implant electrodes

The LFP recordings from the 16 wild-type and 17 KO mice that were previously analyzed [33], were made from a bundle of six 75 μm Formvar-insulated NiCh wire electrodes (California Fine Wire, Grover Beach, CA), staggered by 170 μm, that were stereotaxically implanted under Nembutal anesthesia (60 mg/kg i.p.). The tip was aimed at -1.80 AP, ±1.30 ML, -1.65 DV relative to bregma. The electrodes spanned the dorso-ventral axis of the dorsal hippocampus but the spacing was too great for current-source-density analyses. Reference electrodes were aimed at the cerebellar white matter. For single-unit recordings from 4 wild-type and 3 Fmr1 KO mice, an 8-tetrode, flexDrive (www.open-ephys.org) or a 4-tetrode custom drive was implanted under isoflurane anesthesia (2%, 1 L/min), with the electrodes aimed at the dorsal hippocampus [83], and bone screws, one of which served as a ground electrode. The electrode assemblies were anchored to the skull and bone screws with one of two dental cements (Grip Cement, Dentsply, Milford DE and TEETs Denture Material, Co-oral-ite Dental MMG, Diamond Springs, CA). The mice were allowed at least 1 week to recover before experiments began.

### Electrophysiological Recording

A custom unity-gain buffering preamplifier was connected to the electrode assembly and the electrophysiological signals were galvanically transmitted to the recording system by a light-weight counter-balanced cable. The differential signal from each electrode was low-pass filtered (600 Hz) and digitized at 2 kHz for LFPs and band-pass filtered (500 Hz – 6 kHz) and digitized at 48 kHz for action potentials using dacqUSB (Axona, St. Albans, U.K.). Two millisecond duration tetrode action potential waveforms were collected and stored for offline single unit isolation using custom software (Wclust; see Supplementary information Fig S6). Single unit isolation quality was quantified by computing IsoI_BG_ and IsoI_NN_ [84]. Single units (N = 455) were recorded and analyzed from 6 WT and 3 Fmr1 KO mice. Only 213 single units with both IsoI_BG_ and IsoI_NN_ greater than 4 bits were judged to be well-solated and 163 of these were of sufficiently high quality putative pyramidal cells that express place cell or nonspatial discharge characteristics according to objective criteria (see Supplementary information Fig S6). LFPs were localized as previously described [33] or to CA1 *stratum pyramidale* because they showed LFP activity characteristic of *stratum pyramidale* and were recorded by the same electrode as place cells. The electrode locations were verified histologically at the end of recordings.

### Active Place Avoidance

The active place avoidance task has been described in detail [85, 86] and the behavioral protocol was identical to [33]. Briefly, the mouse’s position was tracked 30 times a second using an overhead camera and a PC-based video tracking system (Tracker, Bio-Signal Group, MA). All sessions lasted 30 minutes. Pretraining on the rotating (1 rpm) arena was followed 24-h afterwards by three training sessions during which the mouse (n = 40) received a mild 0.2 mA, 60 Hz, 500 ms foot shock whenever it entered the shock zone. There was a 2-h rest in the home cage between training sessions. A subset of the mice received conflict training (n = 14) in which the conditions were identical to the training phase, except the shock zone was on the opposite side. The conflict task variant tests cognitive flexibility because the mice should suppress recollection of the initial memories of the location of shock so they can learn and use the new location of shock. Note that because the shock zone is unmarked, the physical conditions of all the sessions are identical except when the mouse is shocked, which is rare; for example, only for 10 s if a mouse receives 20 shocks during a 30-min session.

### Data Analysis

Detection of behavioral events: During the first session of the initial training on the rotating arena, spatial behavior becomes stereotyped, with periods of stillness, when the mouse is passively carried towards the shock zone by the arena’s rotation and periods of movement directed away from the shock zone. This is quantified when angular distance to the shock zone is plotted against time; it reveals a saw-tooth profile (Fig 1B, top). We selected two behavioral events based on the angular distance to the shock zone. The onset of avoidance (blue dots in Fig 1B, top) was defined as local minima in the target angle time series with preceding stillness without entering the shock zone. The second event was an escape (red dots in Fig 1B, top), defined as entrance to the shock zone with preceding stillness. To define stillness, speed was computed using 400 ms long sliding window. Stillness was identified as intervals with speed below 2cm/s. Brief crossings of the stillness threshold for less than 150 ms were not considered departures from stillness. Because some avoidances were preceded with the animal’s initial acceleration towards the shock zone followed by a turn, local minima in the target angle time series occurred during speed above the stillness threshold, we included all avoidances with at least 1 s of stillness in a 3-s window prior to the detected avoidance onset.

Quantifying Vicarious Trial and Error: Vicarious trial and error (VTE) was quantified during SG_dom_, MG_dom_, and randomly-selected events as in prior work [49]. Briefly, the orientation of motion (phi) was first calculated from the position time series as the arctangent of the change in x and y coordinates across 33 ms time steps. The change in phi across 33 ms was then calculated to represent the angular acceleration of motion time series. The absolute value of this time series was summed in 500ms window during an event to estimate the amount of head sweeping in the event.

Preprocessing for LFP recording quality: The LFP data were first processed by a custom artifact rejection algorithm, which marked continuous segments of artifact-free signals. Such segments that were 4 s or longer were used for further analysis. The majority of artifacts were related to the foot shock, specifically signal saturations and slowly changing DC signals as the recording system recovered from the shock artifact. Constant signals close to the maximal voltage of the analog-digital-convertor defined signal saturation. Periods of very low signal difference defined the slowly changing DC signal artifacts. Thresholds for an acceptable signal difference were selected by visual inspection, and used for analysis of the entire dataset. Each artifact segment was extended by 0.25 s on both sides and all artifact-free segments smaller than a 1-s minimum window were also discarded. Each channel in the dataset was processed independently and the algorithm performance was verified by visual inspection.

Detection of oscillatory events: A published algorithm was used to extract oscillatory events from the LFP independently for each recorded channel [31]. In the first step of the algorithm the LFP is transformed into a time-frequency power representation by convolving the LFP signal *S* with a group of complex Morlet wavelets resulting in complex band-specific representations *S_f1_*..*S_fN_*, where *f_1_*..*f_N_* represent frequencies of interest (Fig 1E). Instantaneous normalized power is then obtained by squaring the argument of the band-specific signal representation and dividing it by the square of the standard deviation of the band-specific signal such as 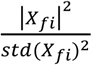. In the next step, oscillatory events are detected as local peaks in the normalized 2-D time-frequency space (Supplementary information Fig S2E). Band-specific oscillation rates are then computed as the number of detected events in a representative frequency bands (30-50 Hz for slow gamma, 70-90 Hz for mid-frequency gamma) with power exceeding a defined threshold (2.5 S.D. of the mean power) per unit of time (Refer to Supplementary Information Fig S3 for the rationale for selecting power thresholds and representative frequency bands).

Calculation of instantaneous SG/MG ratio: First we extracted band-specific oscillatory rates in 1-s long windows advanced by 0.25 s. Next, we smoothed the estimated rates of oscillatory events by 2.5-s long windows and took the ratio of SG (30-50 Hz) to MG (70-90 Hz) oscillatory rates. To obtain the maxima of the SG/MG ratio (SG_dom_ events), we searched for local maxima in the SG/MG series with peak prominence (amplitude difference between the maxima and preceding and following minima) of at least 1 and amplitude above 1 (corresponding to SG > MG). SG/MG minima (MG_dom_ events) were obtained in the same way by finding local maxima in the inverse MG/SG time series using the same prominence and amplitude setting (corresponding to SG < MG).

Bayesian analysis: To obtain estimates of the mouse’s location based on single unit data, we used a published algorithm [38], where the probability of the current location is defined as 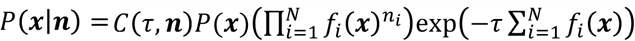, where *C*(*τ*, ***n***) is a normalization factor so that **Σ**_*x*_*P*(***x|n***) = 1, *f_i_*(***x***) are firing rate maps for cells *i..N* obtained either by binning the 2-D space into 32x32 bins or 1-D space (distance to shock zone) into 20 or 12 angular bins, *P*(***x***) is the dwell distribution, *τ* is the length of the time window (500 or 200 ms), *n_i_* is the number of spikes fired by the i-th cell in a given time window and ***x*** is the (*x*,*y*) position of the animal in the 2D analysis or the angular position in the 1D analysis. Only recordings with at least five high quality spatial or non-spatial putative pyramidal cells were analyzed. Time windows with no spikes were excluded from analysis. To obtain the error of the location estimate in 1D, we created a linear error function *E*, which was zero at the observed location and at any other location it corresponded to the distance from the observed location such that *E* = *d*(*x_obs_*, *X_decode_*), where *d* is the distance between the decoded angular bin *X_decode_* and bin, *X_obs_* where the mouse was observed. We then multiplied the location estimate *P*(***x|n***) by the error function and took the average so location errors at the highest probability contributed proportionately more to the resulting error estimate. Considering the entire location estimate *P*(***x|n***) instead of only its maxima lead to a lower overall decoding error and was therefore used throughout this study. The resulting error estimate was z-score normalized to account for absolute differences in the decoded error due to different numbers of putative pyramidal cells in a given recording.

## Statistical analysis

Statistical analyses were performed using JMP version 12 (SAS, Cary, NC) and Matlab 2016b (Mathworks, Natick, MA). Significance was accepted at p < 0.05. Exact p values are reported throughout; p is reported as 0 when p < 10^-125^.

## AUTHOR CONTRIBUTIONS

D.D. analyzed the data, B.R. collected place avoidance and LFP data, F.T.S. and Z.N.T. collected single unit and associated place avoidance and LFP data, A.A.F. supervised research, and D.D. and A.A.F. designed the experiments and wrote the manuscript.

## ACKNOWLEDGMENTS

This work was supported by a grant from the Simons Foundation (294388, A.A.F.), a grant from NIH (R01MH099128) and a CIHR Fellowship to F.T.S.

## SUPPLEMENTAL INFORMATION

### Data and code online repository

To obtain data used for the analyses, please visit https://osf.io/j5baw/?view_only=87b1ff00182a437883da06feda992763

### Basic LFP properties during active avoidance

Local field potentials analyzed during active avoidance training when mice were still (speed < 2 cm/s) and running (speed ≥ 2 cm/s; Fig S1A, B) both expressed peaks in the theta and gamma bands. Power was greater during running than stillness in the 5-10 Hz theta band (t_1,8_ = 4.91, p = 0.001), 30-50 Hz slow gamma band (t_1,8_ = 2.45, p = 0.04) as well as the 70-90 Hz mid-frequency gamma band (t_1,8_ = 6.38, p = 0.0002.

We detected sharp-wave associated ripple events (Carr et al., 2011; Csicsvari et al., 2000; Jackson et al., 2006; O’Neill et al., 2006) using a published algorithm (Eschenko et al., 2008). The rate of ripple events were generally low (Kay et al., 2016) and did not differ during overall stillness or during the stillness that preceded subsequent active avoidance movements (Fig S1C, F_1,28_ = 0.34, p = 0.7).

**Fig S1 related to Fig 1.**
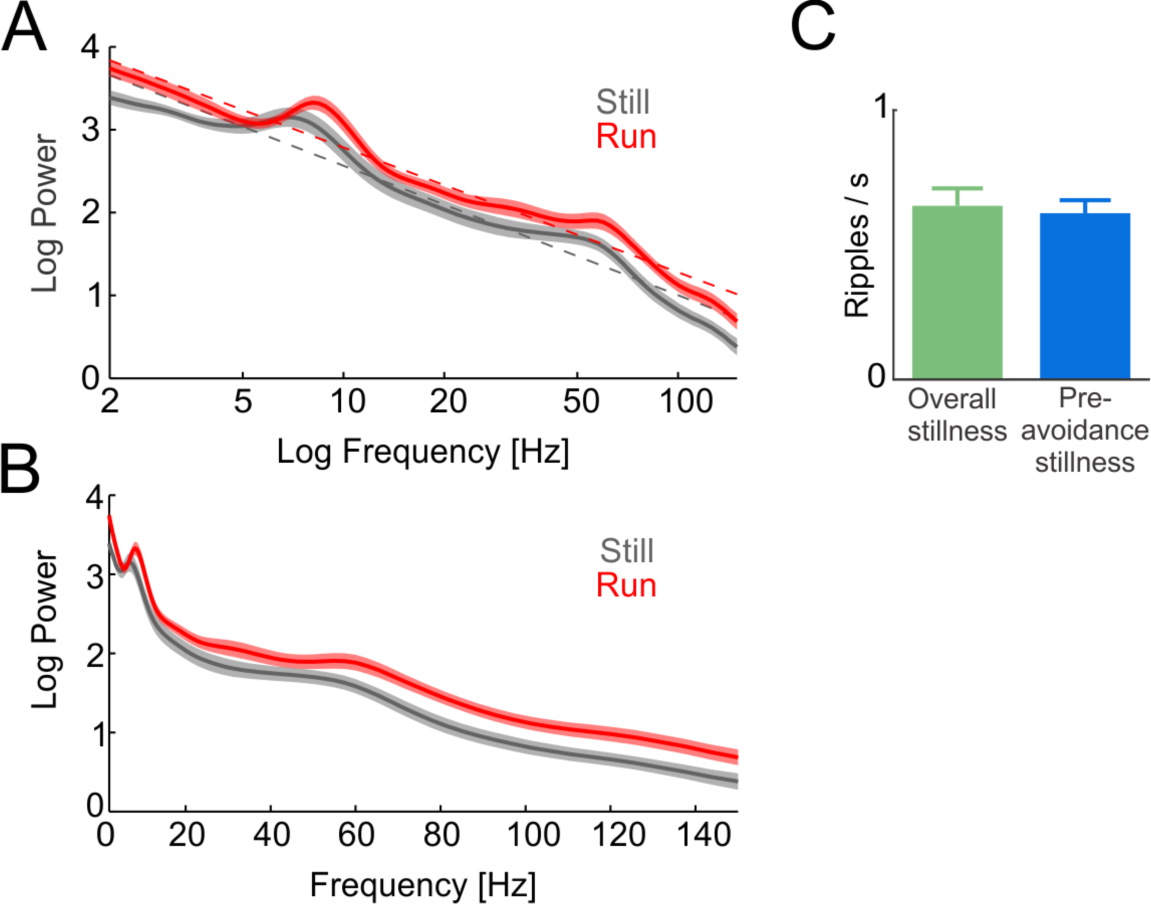
Basic LFP properties during active avoidance. (A) Power spectra with log frequency axis during periods of stillness and running throughout active avoidance training. Dotted lines indicate linear fits to the data. (B) Power spectra with linear frequency axis during periods of stillness and running throughout active avoidance training. (C) Sharp-wave associated ripple rates during pre-avoidance stillness and overall stillness.

**Mixtures of slow**, mid-frequency **and fast gamma oscillations in *stratum pyramidale* of mouse CA1** Concurrent local field potentials (LFPs), reflecting the synchronous synaptic activity at distinct input-defined anatomical locations within the mouse CA1 region of dorsal hippocampus were recorded using 32-ch linear silicon electrode array (Neuronexus, Ann Arbor, MI) in order to demonstrate the presence of three distinct gamma bands within *stratum pyramidale* (sp) as shown previously in rat (Fernandez-Ruiz and Herreras, 2013; Schomburg et al., 2014) and mouse (Lasztoczi and Klausberger, 2014, 2016). LFPs were first localized (Fig S2A) using sharp-wave associated ripples (SWR). The maximum amplitude of the ripple identified *stratum pyramidale*, the maximum amplitude of the sharp wave identified *stratum radiatum*, and the sharp wave reversal identified *stratum lacunosum moleculare*. CSD analysis (Mitzdorf, 1985) was then performed in order to separate individual oscillatory CSD components (Fig S2B). Theta (5-12 Hz) phase was then used to construct theta-averaged CSD power profiles (Fig S2C). While *stratum radiatum* and *stratum lacunosum moleculare* CSDs show the presence of two distinct oscillatory bands (slow gamma 30-60 Hz and mid-frequency gamma 60-120 Hz), *stratum pyramidale* shows three distinct gamma bands (slow gamma 30-60 Hz, mid-frequency gamma 60-90 Hz and fast gamma > 100 Hz). Similar results were obtained when power profiles were constructed using band specific LFPs from *stratum pyramidale* (Fig S2D) during running (speed ≥ 2cm/s) and stillness (speed < 2cm/s). Notably, slow gamma (30-60 Hz) oscillations were distinguished more clearly than in CSD power profiles, while fast gamma (> 100 Hz) was harder to distinguish in LFP power profiles. The ability to detect both slow and mid-frequency gamma oscillations in *stratum pyramidale* is important for two reasons. First, the SG/MG ratio associated with recollection can be sufficiently estimated from the electrodes located in the *stratum pyramidale.* Second, since hippocampal place cell recordings target *stratum pyramidale*, the LFPs recorded together with single units can be used to estimate the SG/MG ratio.

**Fig S2 related to Fig 1.**
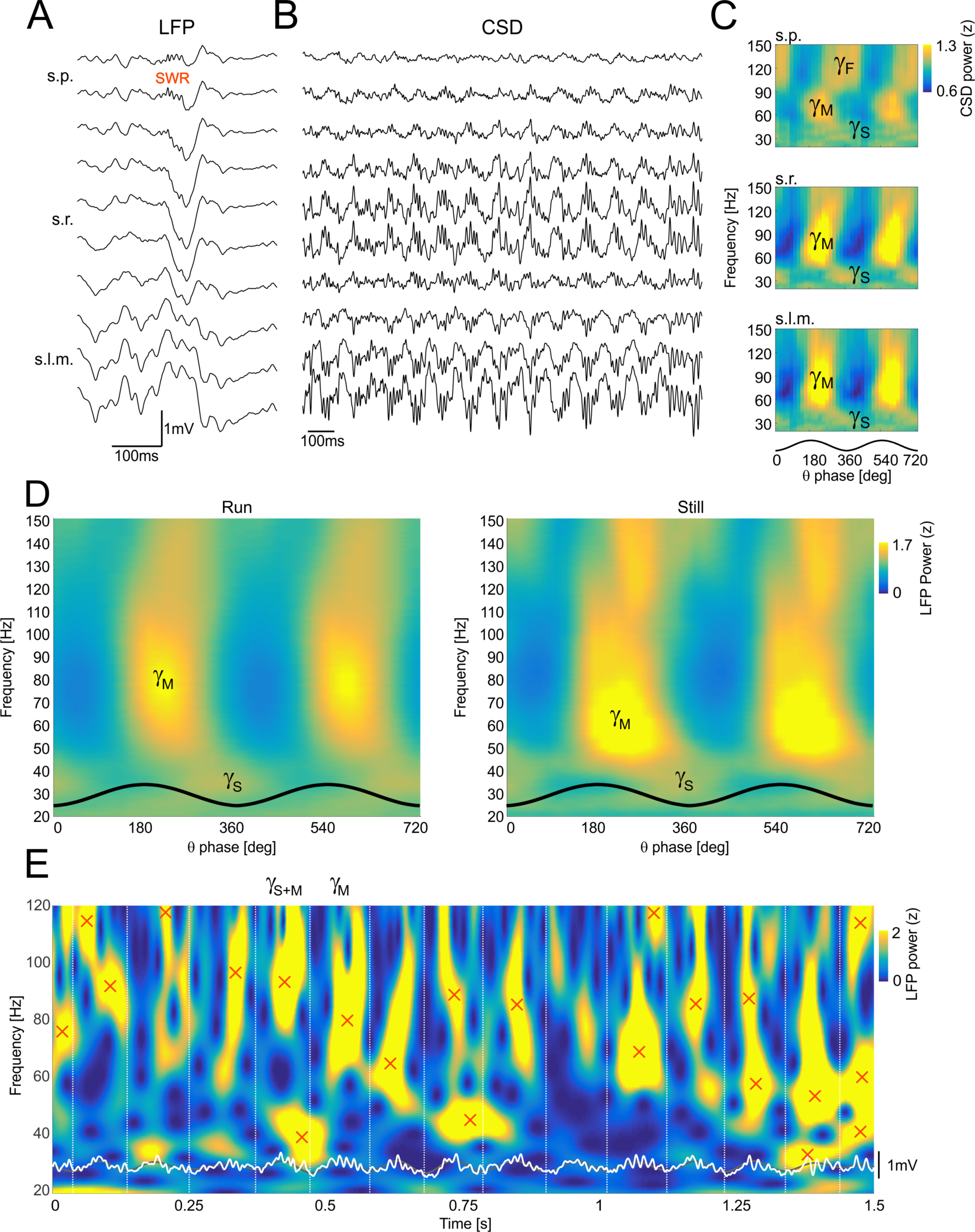
Oscillatory activity in mouse dorsal hippocampus CA1 local field potentials. (A) LFP signals obtained using 32-site linear silicon electrode array during a sharp-wave associated ripple (SWR) s.p – *stratum pyramidale*, s.r. – *stratum radiatum*, s.l.m. – *stratum lacunosum moleculare*. (B) CSD analysis of CA1 LFP to separate individual oscillatory components within CA1. (C) CSD power profiles averaged over theta cycles from s.p., s.r. and s.l.m electrodes. Notice the mixture of three gamma types (slow gamma 30-50 Hz, mid-frequency gamma 60-100 Hz and fast gamma > 100 Hz) at the s.p. electrode. (D) Normalized LFP power from the s.p. electrode averaged across theta cycles for running (speed ≥ 2 cm/s) and stillness (speed < 2 cm/s). (E) Example 1.5 s LFP obtained from the s.p. electrode (white) and its time-frequency representation obtained by wavelet transform. Each frequency band was normalized separately by dividing signal power with signal variance. Notice the presence of mixed states, when slow and mid-frequency gamma oscillations are present in a single theta cycle. Individual theta cycles are marked by vertical lines. Oscillatory events detected as local maxima in time-frequency 2-D space with peak power > 2.5 S.D. are marked with red crosses.

### Mixtures of slow and mid-frequency gamma oscillations during single theta cycles

It was reported that in rat, slow and mid-frequency gamma oscillations tend to occur mutually exclusively during theta cycles (Colgin et al., 2009), but we find instead that slow and mid-frequency gamma oscillations detected as discrete events (Fig S2E) are often mixed in mouse, as reported by others (Lasztoczi and Klausberger, 2016; see Fig S2). It is possible however, that the common, non-exclusive appearance of slow and mid-frequency gamma oscillations in single theta cycles might be specific to mouse.

### Selecting the power threshold for oscillatory events

From the nature of time-frequency representations, when power is reduced, the time-frequency profile loses dominant peaks and instead more low power peaks are detected (Fig S3A). This results in a negative correlation between the number of low power peaks in a given time interval and the total power of all events detected in the interval (Fig S3B). To avoid this bias, we identified an optimal power threshold so that the number of oscillations per unit of time is not influenced by increased counting of low power events. Setting the power threshold to above 2 S.D. from the overall mean band power selects about 35% of all detected events. Setting the power threshold above 3 S.D. selects about 20% of all detected events (Fig S3C). Fig S3D bottom shows the continuous relationship between power threshold applied to detected events in the 30-60 Hz slow and 60-90 Hz mid-frequency gamma range and the ratio of theta cycles classified as having a unique type of oscillation (slow or mid-frequency gamma) and theta cycles classified as having both slow and mid-frequency gamma oscillations. We only included periods when the mouse was running (speed ≥ 2 cm/s) and only included theta cycles with periods between 83 and 250 ms corresponding to frequencies between 4 and 12 Hz. Predictably, low power thresholds lead to high numbers of detected events but a low ratio of single types of event (slow or mid-frequency gamma) present in a given theta cycle (Fig S3D top, left; theta cycles only with slow gamma ‘S’, mid-frequency gamma ‘M’, mixture of slow and mid-frequency gamma ‘S/M’ and no detected oscillations ‘Ø’). In contrast, high power thresholds lead to a low number of detected events and a high prevalence of single gamma-type theta cycles (Fig S3D top, right).

Because of the continuous relationship between power threshold and the ability to classify a theta cycle as one with just slow or mid-frequency gamma, we used the ratio of slow to mid-frequency gamma events per time period throughout this report.

### Identifying a power threshold and frequency bands of interest for slow and medium gamma event rates

To investigate the apparent asymmetry in the relative declines of slow and mid-frequency gamma events that was observed in the wavelet spectrum prior to avoidance (Fig 1E in the main text), we first identified the specific frequency bands of interest (Fig S3E). We computed the rate of oscillatory events in 1-s long intervals that advanced by 250 ms. The event rates around the avoidance onset were averaged across 20 Hz-wide bands between 20 and 110 Hz and for power thresholds z ≥ 1, 2, 2.5 and 3 during the third training session. We then compared these profiles across all bands to look for similarities. The 20-40 Hz band followed similar patterns of activity as the other two slow gamma bands (30-50 Hz and 40-60 Hz), but its average event rates were generally low so we excluded this band from further analysis. The 30-50 Hz and 40-60 Hz slow gamma bands were distinct from bands in the mid-frequency gamma range (60-110 Hz) for power thresholds above z ≥ 1, because the attenuation of slow gamma was reduced prior to avoidance onset, in fact, there were peaks of increased slow gamma rates 2-4 s before the avoidance initiated and there was an earlier increase of the slow gamma event rate at avoidance onset compared to a delayed increase in the rate of mid-frequency gamma events. We selected the 30-50 Hz band to represent slow gamma (SG) oscillations and the 70-90 Hz band to define mid-frequency gamma (MG) oscillations. In all subsequent analyses we only included events that were 2.5 S.D. or greater than the average power in a given frequency band.

**Fig S3 related to Fig 1.**
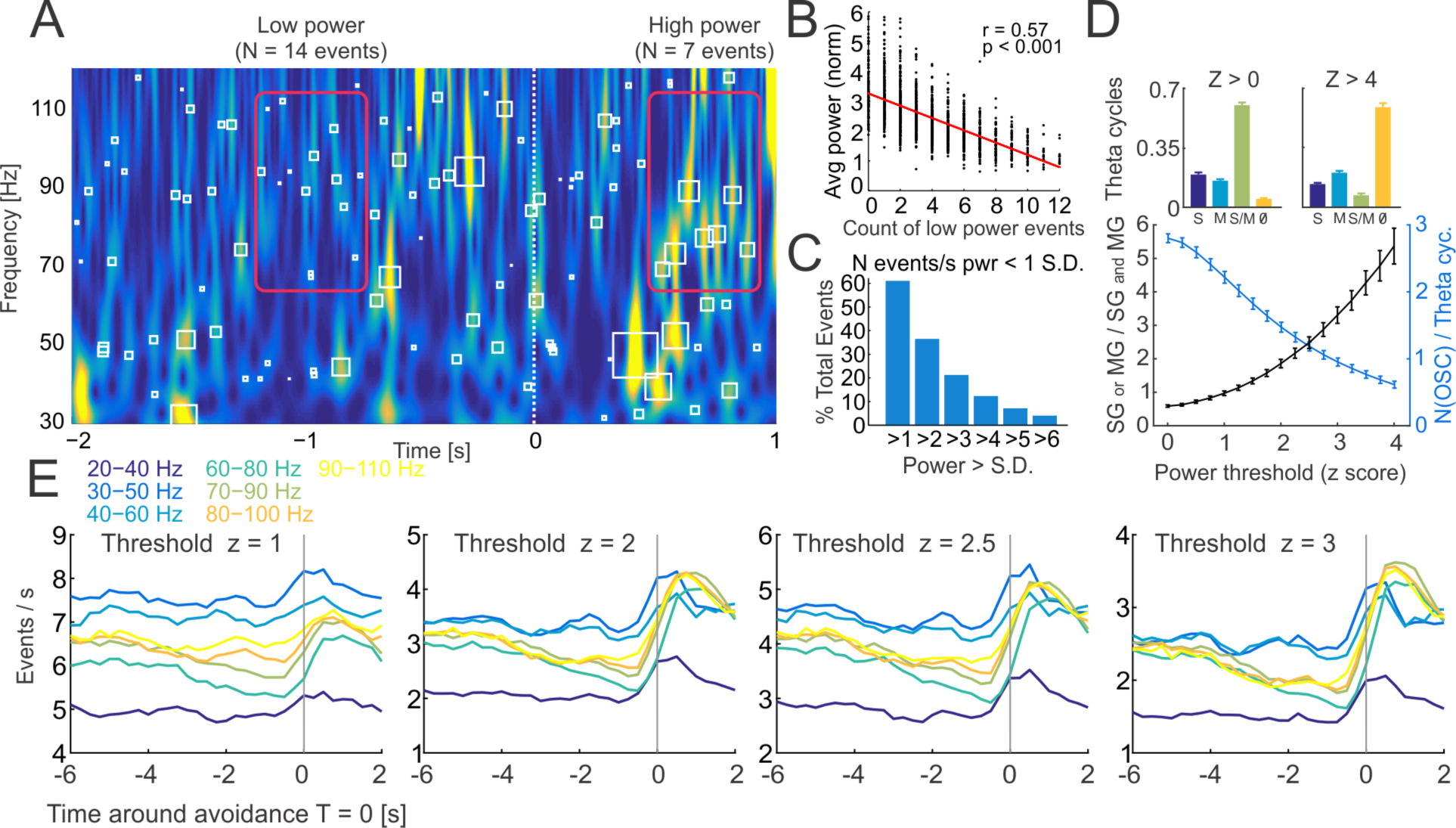
Selecting the threshold for oscillatory events and selecting the oscillation frequencies to represent slow and mid-frequency gamma. (A) Example time-frequency representation of LFP before and after avoidance. Power in the mid-frequency gamma range during stillness prior to avoidance is typically attenuated, leading to a higher number of detected low power events (N = 13, red rectangle around T = -1 s). Power in both slow and mid-frequency gamma ranges is typically increased during running away from the shock zone leading to a higher number of detected high power events (N = 7, red rectangle around T = +0.75 s). (B) Average normalized power in a 1-s interval is negatively correlated with the number of low power events (z < 1) in the interval. (C) The proportion of detected events after applying different power thresholds. (D) Top: ratio of detected theta cycles with only slow gamma (S), mid-frequency gamma (M), mixture of slow and midfrequency gamma (S/M) and no detected oscillations (Ø) for power threshold z > 0 (left) and z > 4 (right). Bottom: Relationship between the power threshold and the ratio of theta cycles with a single type of oscillation (SG or MG) and the ratio of theta cycles with mixed oscillations (SG and MG; black) plotted together with the number of supra-threshold oscillations per theta cycle (blue). (E) The average oscillation rates for 20-Hz wide bands covering the 20-110 Hz frequency range around avoidance onset (T=0) for power thresholds z ≥ 1, 2, 2.5 and 3. Only the average profiles are included for clarity.

**Fig S4 related to Fig 3.**
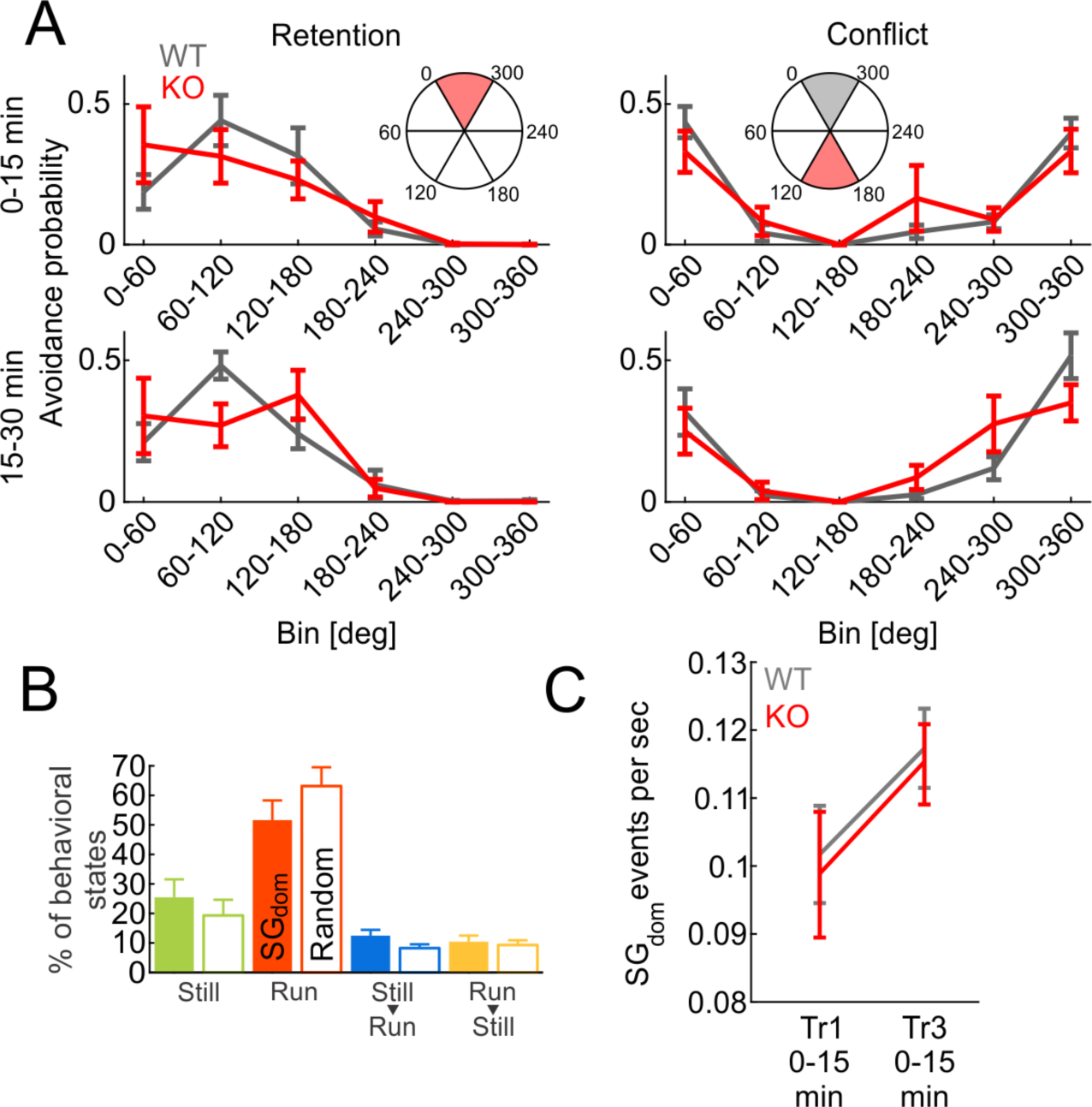
Avoidance histograms during Retention and Conflict sessions, probability of behavioral states during SGdom and random events in pretraining sessions and SG_dom_ event rates across training sessions. (A) Locations of avoidances across six 60° sectors during the first and second halves of the memory retention and conflict sessions. Avoidance profiles analyzed by two-way ANOVA with repeated measures, only the second half of the Conflict session (15-30 minutes; Fig S4A bottom, right) shows a significant genotype x bin interaction (F_5,8_ = 4.56, p = 0.03) as Fmr1-KO animals display stronger avoidance of locations associated with the initial shock zone (0-60 deg, 300-360 deg). (B) Proportions of different behavioral events detected during SG_dom_ events during pretraining sessions before ever experiencing shock (filled bars) compared to randomly-selected events (empty bars; Comparisons of SG_dom_ to Random Still: 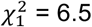, p = 0.16; Run: 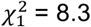, p = 0.08; Stil→Run: 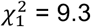, p = 0.05; Run→Still: 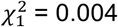, p = 0.99). (C) Average SG_dom_ rates across initial 15 minutes of first and last training session (two-way ANOVA with repeated measures genotype x trial: genotype: F_1,13_ = 0.30, p = 0.59; trial: F_1,13_ = 6.91, p = 0.02; genotype x trial: F_1,13_ = 0.04, p = 0.85).

### Relationship between decoding accuracy, firing rate and speed

**Fig S5 related to Fig 4.**
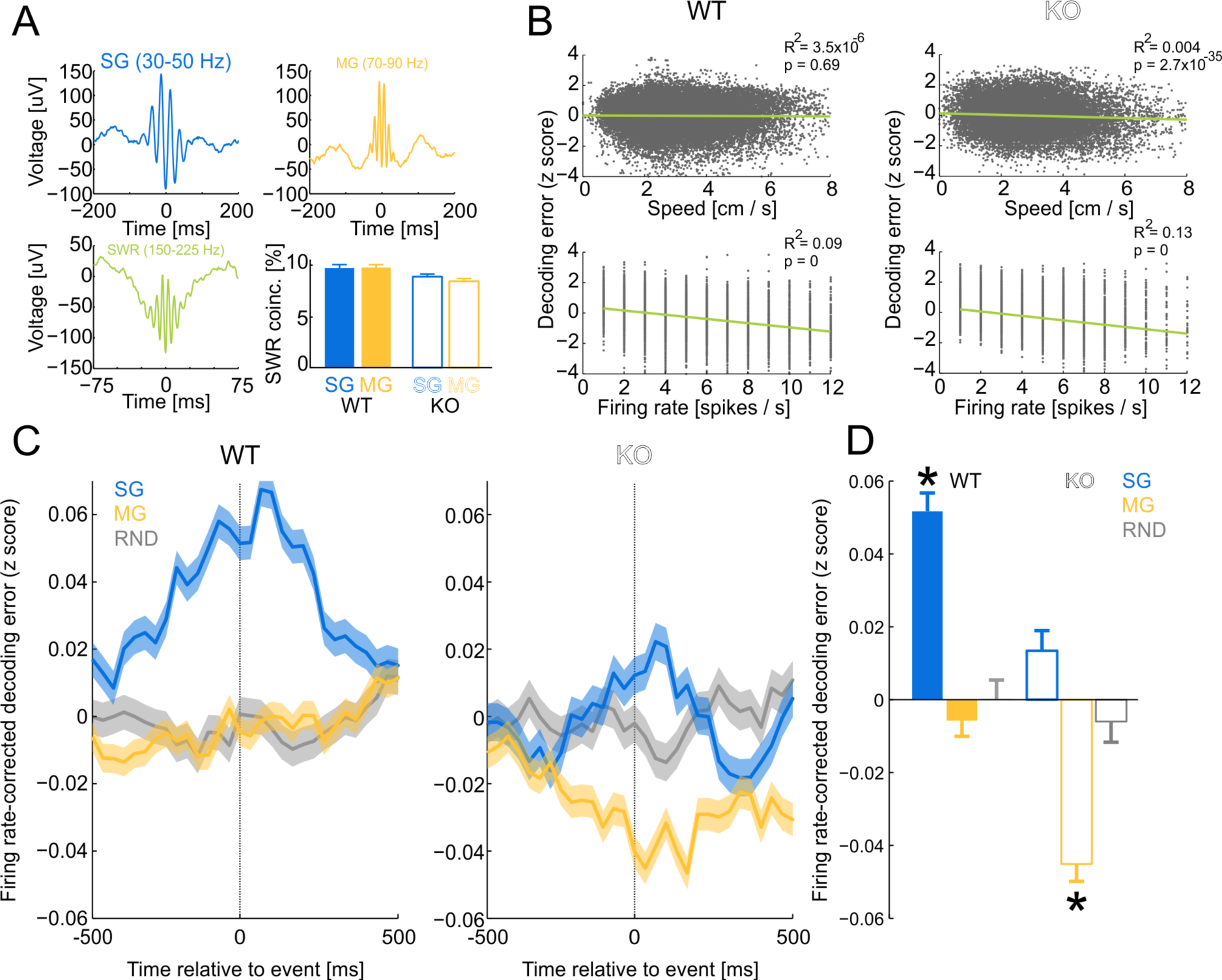
Non-local place representations in putative pyramidal cell ensemble discharge during slow gamma oscillations without concurrent mid-frequency gamma oscillations and sharp-wave associated ripples in wild-type and Fmr1-KO mice. (A) Average voltage traces during isolated slow gamma (SG), mid-frequency gamma (MG) and sharp-wave associated ripple (SWR) events. Percentage of (non-isolated) SG and MG events that coincide with SWR events. (B) Relationships between location decoding error (smaller error = more accurate) and running speed (top) and ensemble firing rate (bottom). (C) Bayesian decoding error during slow gamma oscillations that are not accompanied by mid-frequency gamma oscillations or SWRs (blue), mid-frequency gamma oscillations that are not accompanied by slow gamma oscillations or SWRs (yellow) and random events, which are not accompanied by SWRs (gray) in WT and KO mice, corrected for firing-rate bias of the decoding. (D) Summary of decoding error during isolated slow and mid-frequency gamma oscillations and random events. *p < 0.05 relative to random events.

**Fig S6 related to Fig 4.**
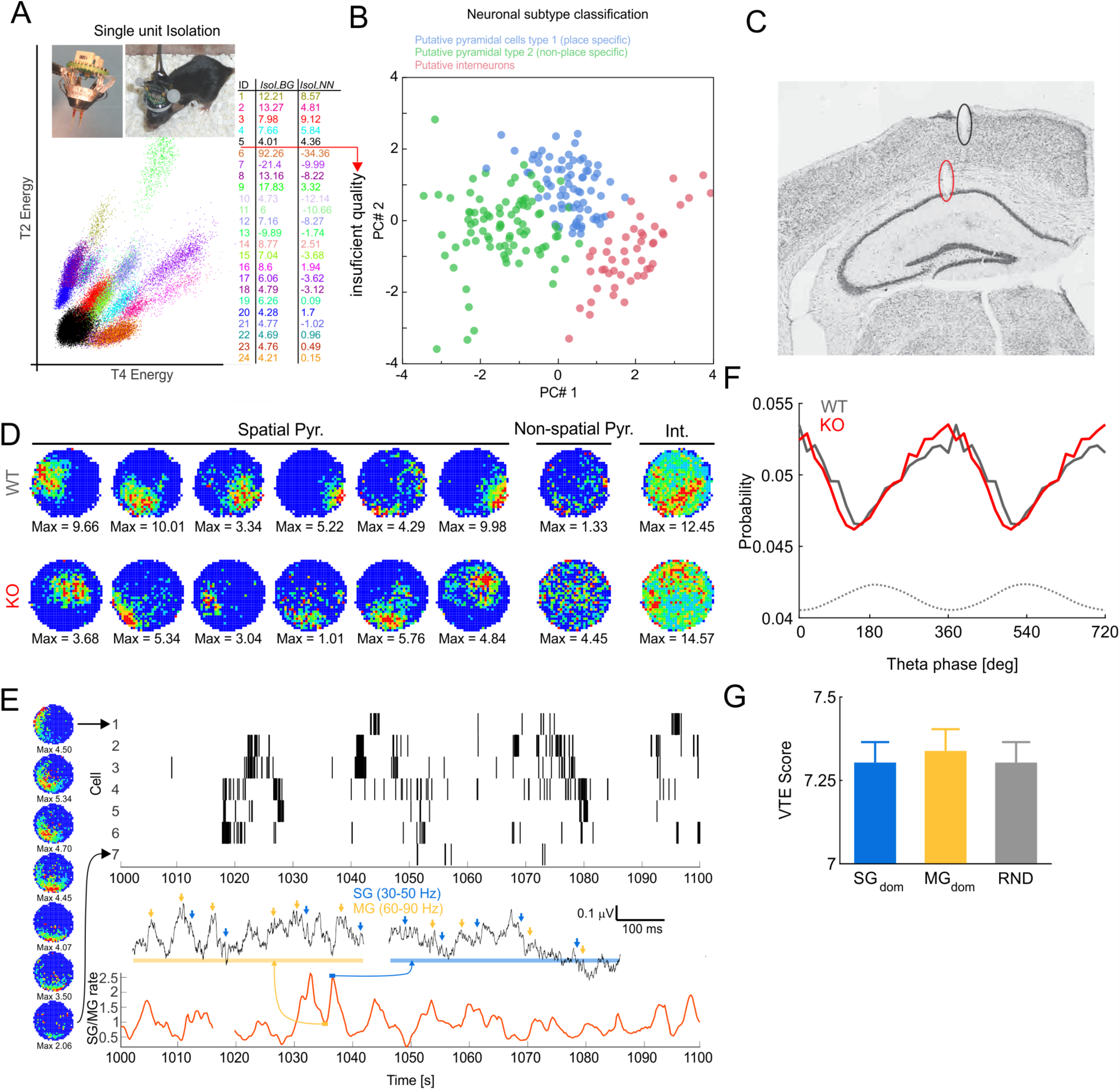
Single unit recordings and selection of the pyramidal cell data set. (A) Example of single unit isolation, and the Open Ephys microdrive (left inset) and an implanted mouse (right inset). The inset table lists isolated units with their corresponding *IsoI_BG_* and *IsoI_NN_* values used for selecting units with sufficient isolation quality. Colors in the table correspond to clusters on the left. Only units with both quality measures > 4.0 were analyzed further. (B) Neuronal subtype classification into three subtypes representing putative pyramidal cells with spatial specificity (type 1; blue), putative pyramidal cells without spatial specificity (type 2; green) and putative interneurons (red). Each dot represents a single well-isolated unit. Plot in 2-D principal component space computed from the original 7-D feature space that describes each unit. These features are: the largest spike’s width, firing rate, proportion of active pixels, firing-rate map coherence, firing-rate map information content, peak ISI and proportion of spikes in a burst (≤10ms ISI). (C) Histology showing electrode placement in CA1. Red and black ellipses mark tip of tetrode and point of entering cortex respectively. (D) Example spatial putative pyramidal cells, non-spatial putative pyramidal cells and putative interneurons from example wild-type (top row) and Fmr1-KO (bottom row) animals. (E) A 7-cell ensemble of spatially-tuned putative pyramidal cells with their corresponding firing rate maps (left) and raster plots of firing (top, right) during 100 s. The corresponding SG/MG ratio is shown in red (bottom, right) with LFP waveforms around the SG/MG maxima and minima with identified SG (blue arrows) and MG (yellow arrows) oscillatory events. (F) Theta (8 Hz) phase preference of spatially-tuned putative pyramidal cell discharge for wild-type (gray) and Fmr1-KO (red) mice. (G) Vicarious trial-and-error score computed for SG_dom_, MG_dom_ and random events.

**Fig S7 related to Video S1.**
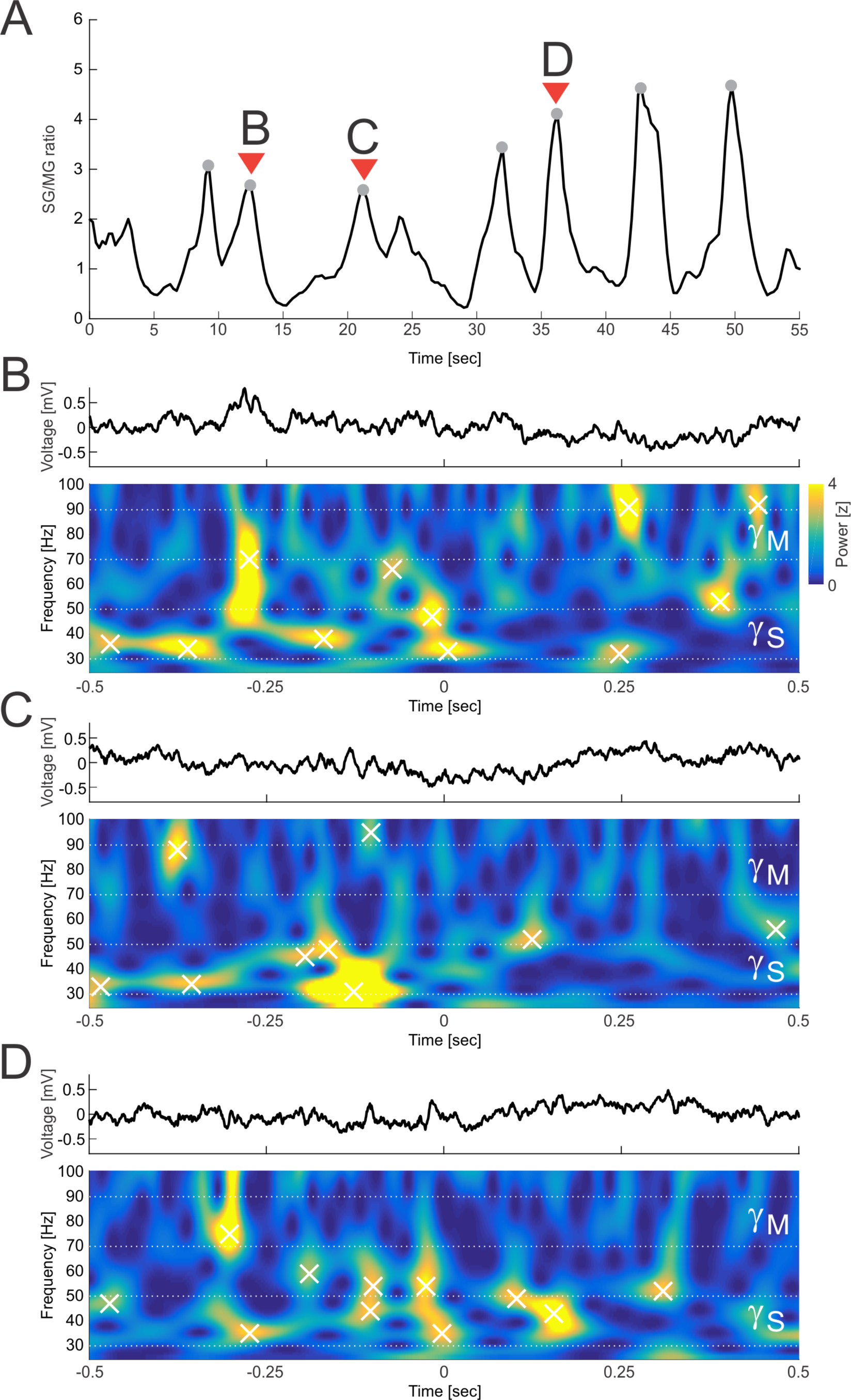
Raw LFP and wavelet spectrum examples of identified SG_dom_ episodes. (A) SG/MG ratio computed over a 55-s long time period with identified SG_dom_ peaks (gray circles) and a selected subset of SG_dom_ peaks used for LFP and wavelet spectrum extraction in the examples below. (BD) LFP (top) and wavelet spectrum (bottom) of 1-s long segments centered on SG_dom_ peaks. Power peaks with z-score power > 2.5 were marked by white cross.

**Video S1 related to Fig 2. Dynamical schematic representation of SG_dom_-identified recollection**. Real-time slow (30-50 Hz; low pitch) and mid-frequency (70-90 Hz; high pitch) gamma event replay represented as an audio track. Notice how both SG and MG events co-occur but that SG events can dominate. Top, left: Polar coordinate of the mouse in a 55-s long window (0 degrees corresponding to the leading edge of the shock zone). The current time in the video is indicated by the vertical line. Middle, left: Slow and mid-frequency gamma rates. Bottom, left: SG/MG ratio. Top, right: Location of the mouse in the place avoidance environment with the shock zone shown in red. The current location is indicated in black and the prior 6 s (180 samples) of locations are shown in gray-scale from black to white. Bottom, right: time-frequency representation of the LFP obtained from *stratum pyramidale* in the 25-100 Hz range during 5-s long windows centered around the current time (vertical line on the left). The video uses the same data that are presented in Fig 2 and Fig S7.

